# Selective autophagy of whole micronuclei suppresses chromosomal instability

**DOI:** 10.64898/2026.04.03.716211

**Authors:** Nikolaus A. Watson, Mattia Melli, Marco R. Cosenza, Viola Oorschot, Lisa B. Frankel, Jan O. Korbel, Francesco Cecconi

## Abstract

Chromosomal instability (CIN) arising from mitotic errors generates pervasive structural and numerical chromosome alterations fueling cancer evolution^1–3^. Entrapment of missegregated chromosomes within micronuclei exacerbates CIN by fostering repeated rounds of aberrant mitotic segregation^4–6^ and, following micronucleus rupture, promoting catastrophic chromosomal rearrangement processes such as chromothripsis^5–9^. Whether cellular mechanisms exist to restrain micronucleus-driven CIN has remained unclear. Developing a live-cell chromatin acidification sensor, we tracked micronuclei from genesis through subsequent cell cycles and observed frequent whole-micronucleus capture and acidification via the autophagy pathway. Autophagic targeting is selective for micronuclei with nuclear envelope defects seeded at mitotic exit. Our data indicate that these defects drive progressive loss of chromatin–nuclear envelope tethering, producing a mechanically altered state that is recognised by the autophagic machinery. Autophagy and rupture represent distinct micronuclear fates with opposing genomic consequences. Single-cell sequencing of fate-matched cells demonstrates complete digestion of the chromosomal contents of autophagy-targeted micronuclei, a process we term chromophagy (chromosome-autophagy). By eliminating micronuclei, chromophagy promotes chromosomal loss and arrests the intergenerational transmission of missegregated chromosomal material. This constrains the mutational consequences of micronucleation and suppresses micronucleus-mediated CIN.

## Introduction

Chromosomal instability (CIN) is an enabling cancer hallmark^10^, driving metastasis, therapy resistance, and rapid karyotype evolution through ongoing structural and numerical chromosome alterations^1–3^. Cells exhibiting CIN incur chronic errors in cell division, resulting in the sequestration of mis-segregated chromosomes within micronuclei^11,12^. These aberrant nuclear compartments often emerge recurrently across consecutive cell cycles^4–6^. They frequently rupture during interphase and undergo defective DNA replication and repair, producing structurally altered and compromised chromosomes that are vulnerable to repeated mis-segregation and fragmentation^5,6^. Chromosomal fragments entrapped within micronuclei have been demonstrated to undergo catastrophic rearrangement processes, including breakage–fusion–bridge cycles and chromothripsis^5–7,13–15^. These processes are capable of rapidly reconfiguring karyotype structure, and can give rise to extrachromosomal DNA in cancer^2^. As such, micronuclei act as engines of CIN, fueling accelerated somatic karyotype remodelling^2,16,17^. It remains unclear whether cells deploy mechanisms capable of constraining this escalating micronucleus-mediated genetic instability.

Autophagy is a conserved degradative process that maintains energy balance and cellular quality control by sequestering cytoplasmic material, including damaged organelles, within double-membrane vesicles for lysosomal degradation^18^. Several key members of the autophagy pathway have been reported to occasionally localize to micronuclei^19^, and pathway disruption has been reported either to impact the rupture status^20^ or, in some studies, the frequency of micronuclei^21–23^.

To investigate the consequences of autophagic targeting of micronuclei, we developed a live-cell chromatin acidification sensor and tracked micronuclei from their genesis through subsequent cell cycles. We find that a subset of micronuclei undergo autophagy-mediated lysosomal acidification across multiple human cell lines. This process involves whole-micronucleus capture within large autophagosomal structures followed by lysosomal fusion and acidification, and it selectively targets micronuclei derived from lagging chromosomes that exhibit defective nuclear-envelope assembly. Our data support a model in which these micronuclei incur a progressive loss of chromatin–nuclear envelope tethering, generating a mechanically distinct state recognized by the autophagic machinery. By coupling live-cell imaging with homolog-resolved template strand genomic sequencing in sister cells, we show that micronucleus-autophagy removes entrapped chromosomes from the genome, limiting the propagation of genomic material prone to undergo further chromosomal rearrangements. Micronucleus-autophagy thus offers a potential explanation for the previously described selective loss of whole chromosomes in cancer genomes^6,24,25^. In this manner, it constrains micronucleus driven CIN, and arrests cycles of chromosome mis-segregation and fragmentation which fuel accelerated karyotype evolution in cancer.

## Results

### Whole-micronucleus autophagy in human cells

To assess the potential consequences of micronuclei-localised autophagy-pathway activity, we developed a live-cell chromatin acidification reporter consisting of histone H2B fused to the pH-sensitive fluorophore mKeima (H2B-mKeima) **(Fig 1a)**^26^. We induced H2B-mKeima expression in transformed (U2OS and HeLa) and non-transformed (hTERT RPE-1) human cell lines, and monitored the fate of spontaneously arising micronuclei from genesis through the subsequent cell cycle by live-cell imaging. In all cell lines tested, ∼10-20% of micronuclei exhibit a sudden shift in fluorescence signal intensity from neutral to acidic mKeima channels, indicating chromatin acidification **(Fig 1b, c)**. Micronuclei induced using small molecule inhibitors, known to compromise faithful mitosis^13,14,27^, also undergo acidification at similar rates, with the notable exception of low dose taxol treatment, which produces large micronuclei, which do not undergo acidification **(Fig 1d).** We find that a widely employed nocodazole release procedure^12,13^ results in particularly high acidification rates (around 30%) **(Fig 1d)**, and we therefore utilised this procedure for micronuclei induction in subsequent experiments, unless otherwise stated.

**Figure 1:**
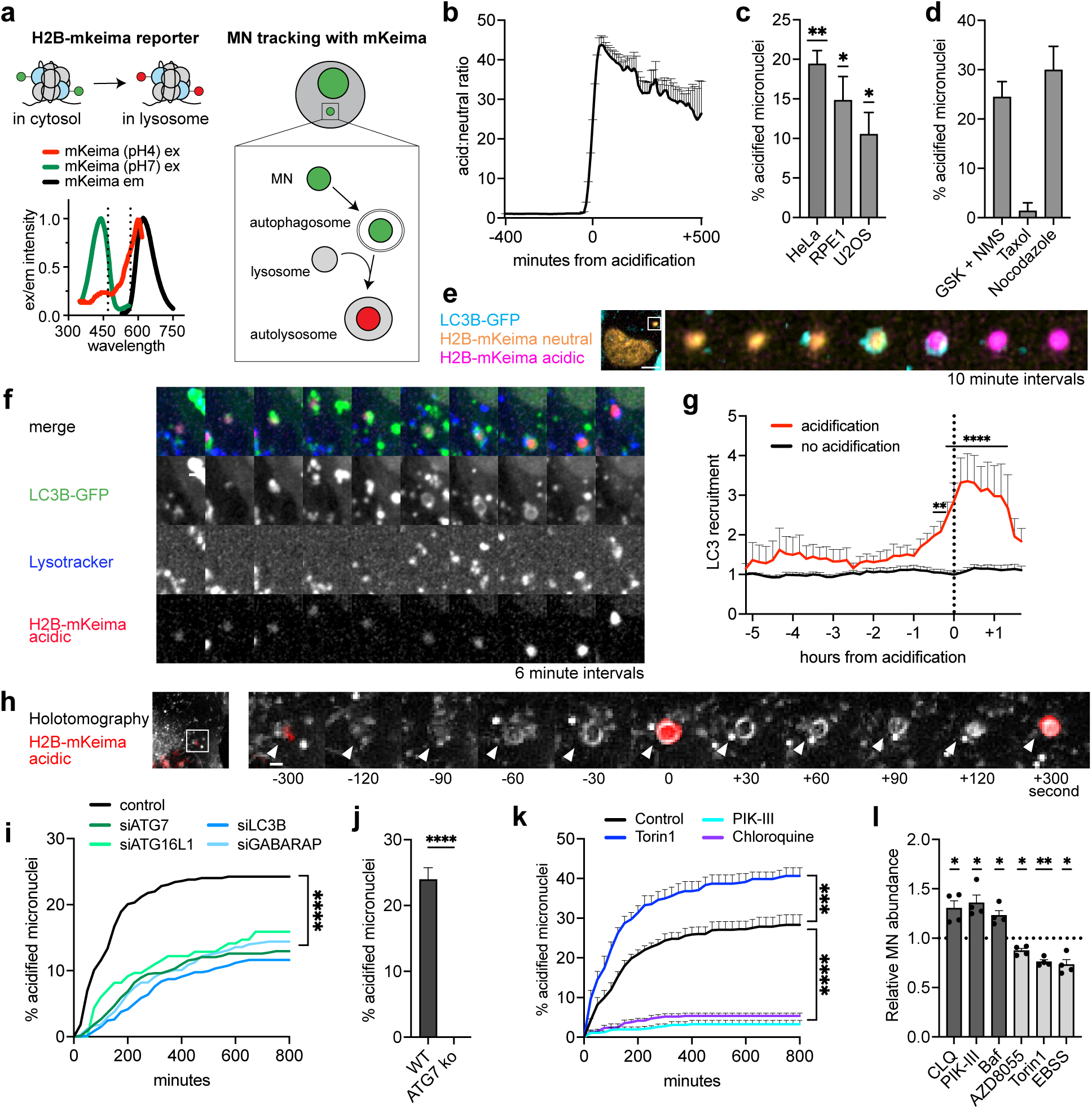
Autophagic capture and acidification of whole micronuclei. **a,** Schematic of the H2B-mKeima chromatin acidification reporter. Primary excitation shifts from 488nm to 561nm laser upon acidification **b,** H2B-mKeima acidic to neutral signal ratio change in micronuclei upon acidification (n=30MN). **c,** Acidification rate of spontaneously arising MN in different cell lines (n between 20 and 50 MN per cell line per experiment, 3 separate experiments). **d,** Acidification rate of MN induced with different methods (n≥100MN per experiment, 3 separate experiments). **e,** Montage showing MN targeted by LC3B, engulfed in an autophagosome-like structure, followed by H2B-mKeima acidification (scale bar = 5μm). **f,** Montage showing MN capture within LC3B positive autophagosome, followed by lysosome fusion and reporter acidification (scale bar = 3μm). **g,** Quantification of LC3 recruitment on MN before reporter acidification (n=20 MN per fate). **h,** Holotomography montage showing membrane recruitment around MN before acidification of the reporter (scale bar = 2μm). **i,** MN acidification rates over the course of 14h after nocodazole shake-off upon silencing of main autophagic components (n≥100MN per experiment, 4 separate experiments). **j,** MN acidification rate in ATG7ko cells (n≥100MN per experiment, 4 different clones). **k,** MN acidification rates in cells treated with autophagy modulators (n≥100MN per experiment, 4 separate experiments). **l,** MN abundance in cells treated for 16h with autophagy modulators after nocodazole shake-off (n≥500 cells per experiment, 4 separate experiments). Data are shown as mean ± sem, and were analysed using a one-sample t test (c, l), two-way ANOVA (g, i, k), unpaired t-test (j) (* = p ≤ 0.05, ** = p ≤ 0.01, *** = p ≤ 0.001, **** = p ≤ 0.0001)

The acidification signal we observed affected the entire chromatin content of micronuclei, suggesting whole-micronuclei autophagosomal capture, followed by lysosomal fusion. To verify this, we employed multichannel live-cell imaging to monitor the localization and dynamics of autophagosomes (LC3B-GFP) and lysosomes (SiR-Lysosome) relative to acidified micronuclei. We find that acidification events are consistently preceded by stepwise targeting and capture of entire micronuclei within LC3B-GFP positive vesicles **(Fig 1e-g, S1a),** followed by fusion of these structures with lysosomes (**Fig 1f).** Using holotomography imaging, which allows label-free visualization of membrane structure, the formation of an autophagosome-like structure around the micronucleus is evident before acidification of the mKeima reporter **(Fig 1h)**. In addition, we observe similar results using H2B-GFP-mCherry, an alternative acidification reporter adapted from constructs commonly used in selective autophagy research^28^ **(Fig S1b).**

To further verify the dependence of these events on the autophagy machinery, we monitored micronuclei acidification following siRNA-mediated depletion of key autophagy factors^18^. We observe that micronuclei acidification rates are reduced to approximately 50% of control levels following siRNA-mediated depletion of ATG7, ATG16L1, LC3B or GABARAP **(Fig 1i, S1c)**, consistent with a central role for both the core autophagy machinery and the autophagy cargo-adapter recruiters (mATG8) in delivering micronuclei to lysosomes. We then tested the role of the different ATG8 family proteins and found that both LC3s and GABARAPs are involved in the process **(S1d,e)**. We also generated clonal ATG7KO cell lines lacking functional autophagy, observing complete abrogation of micronuclei acidification **(Fig 1j, S1f)**. To rule out indirect effects on the micronucleus population we explored post-mitotic, pharmacological inhibition or induction of autophagy. Inhibition of the lipid kinase VPS34 (PIK-III), which plays an essential role in the early steps of autophagy^29,30^, or inhibition of lysosome function (chloroquine), completely blocks micronuclei acidification **(Fig 1k)**. Conversely, induction of macroautophagy through inhibition of mTOR (Torin1) increases total micronuclei acidification rates substantially **(Fig 1k)**. In line with this, autophagy inhibitors increase micronuclei numbers, while autophagy inducers reduce the amount of micronuclei **(Fig 1l)**. We conclude that a substantial proportion of micronuclei are delivered to lysosomes via the autophagy pathway in human cells.

### Rupture and autophagy are alternative micronucleus fates

Micronuclei often incur irreversible nuclear envelope rupture during interphase^11,12,31^. This event exposes micronuclei entrapped chromosomal material to the cytosol, where it accumulates DNA damage and fragmentation, thereby initiating localized genomic catastrophes such as chromothripsis^5,7,14,32–34^. We reasoned that loss of nuclear membrane compartmentalisation could have a substantial impact on the likelihood of a micronucleus to undergo autophagic capture and acidification, and that determining the relationship between acidification and rupture events could provide important insights into the mechanisms and consequences of micronucleus-autophagy.

To assess both rupture and acidification events in live cells we constructed cell lines expressing reporters of chromatin acidification (H2B-mKeima), nuclear compartmentalisation (HaloTag-NLS), and cytosolic DNA exposure (BAF-GFP). Consistent with previous reports^35^, we find that rupture events are characterised by a sudden loss of nuclear compartmentalisation, and massive recruitment of BAF **(Fig 2a,b, Fig S2a,b)**. Strikingly, only a tiny fraction (<2%) of micronuclei subject to rupture subsequently undergo acidification **(Fig 2c),** indicating that micronucleus rupture may preclude micronucleus-autophagy. In line with this, almost all micronuclei which undergo acidification maintain nuclear compartmentalisation up to and after acidification and do not show BAF recruitment **(Fig 2a-d).** Furthermore, holotomography imaging shows that micronuclei undergoing rupture do not recruit membranes in the same way those destined for autophagy do **(Fig S2c)**. We therefore conclude that rupture and autophagy represent distinct and essentially non-overlapping micronucleus fates.

**Figure 2:**
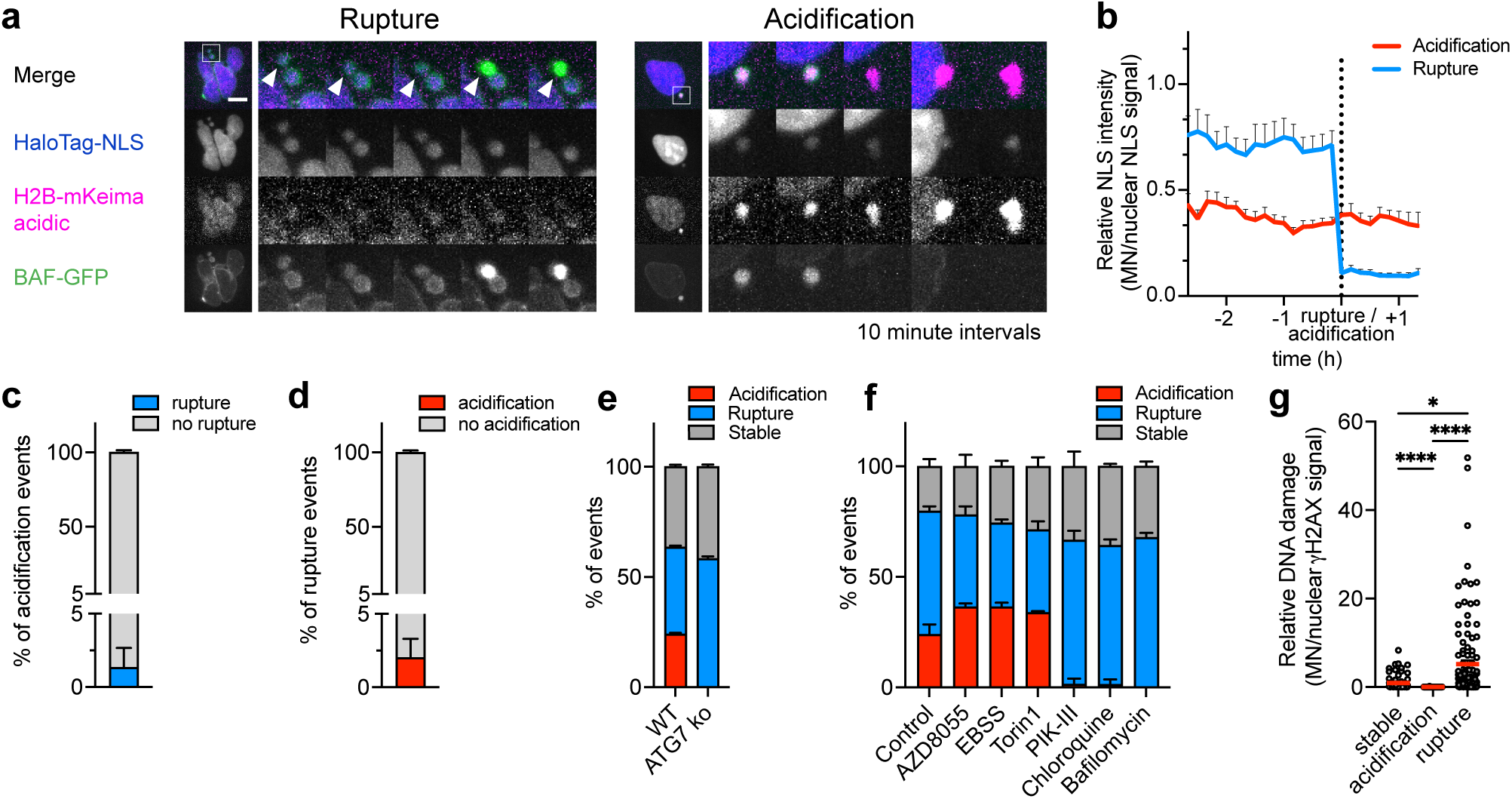
Micronucleus autophagy limits micronuclear rupture and interphase chromatin fragmentation. a, Representative images showing a rupture event (left, indicated with arrow) and an acidification event (right) (scale bar = 10μm). b, Quantification of nuclear compartmentalization signal in micronuclei (HaloTag-NLS) before and after micronucleus (MN) rupture or acidification (n=20MN per fate, 2 separate experiments). c, Quantification of acidification events with or without prior membrane rupture (n≥100MN per experiment, 5 separate experiments). d, Quantification of rupture events with or without prior micronuclear acidification (n≥100MN per experiment, 5 separate experiments). e, Quantification of MN fate distribution in ATG7ko cells (n≥ 100MN per experiment, 2 clones, 2 separate experiments). f, Quantification of MN fate distribution upon autophagy modulation (n ≥100MN per experiment, 3 separate experiments). g, Quantification of MN DNA damage (relative to the primary nucleus) for different MN fates (n between 57 and 116 individual MN, 3 separate experiments). Data are shown as mean ± sem, and were analysed using a one-way ANOVA (g) (* = p ≤ 0.05, **** = p ≤ 0.0001)

As micronucleus autophagy and rupture fates are largely non-overlapping, we next asked whether altering autophagy rates would reciprocally affect rupture frequencies. In support of this notion, rupture rates are significantly elevated in ATG7KO clones without functional autophagy pathway activity, compared to the parental cell line **(Fig 2e)**. To rule out any indirect effects on the composition of the micronucleus population and ensuing fates, we next employed small molecule inhibitors to perturb autophagy pathway activity only *after* mitotic exit. Acute macroautophagy stimulation through Torin1 inhibition (AZD8055, Torin1) or starvation (EBSS) boosts micronucleus acidification rates while markedly reducing rupture **(Fig 2f)**. By comparison, inhibition of VPS34 (PIK-III) or lysosome function (Chloroquine, Bafilomycin) almost completely abolishes acidification, while significantly elevating micronucleus rupture frequencies **(Fig 2f)**. These data further bolster the conclusion that micronucleus-autophagy and rupture represent distinct, non-overlapping, micronucleus fates and indicate that there is competition between these fates in determining micronucleus outcomes.

To compare the impact of these distinct fates on interphase chromosome integrity, we stained live-imaged micronucleated cells with yH2AX to detect DNA double-strand breaks. As has been reported previously^12,13^, a large proportion of post-rupture micronuclei exhibit extensive DNA damage. Conversely, γH2AX was undetectable in all post-acidification micronuclei examined **(Fig 2g)**. In line with this, γH2AX levels were substantially reduced in the micronuclei of cells released into an autophagy stimulator (AZD8055), and increased following release into an autophagy inhibitor (PIK-III) **(Fig S2d)**. These data further confirm that rupture and micronucleus-autophagy are distinct fates, and indicate that they have an inverse impact on the integrity of micronucleated chromosomal material.

### Mitotic exit dynamics predict micronucleus fate

The specificity of micronucleus targeting observed by live imaging (see **Fig 1e)** led us to consider whether micronucleus-autophagy may be a selective process. Selective forms of autophagy often act on damaged or otherwise functionally compromised substrates^36^. Chromosomes persisting in a lagging position during nuclear envelope reassembly often give rise to micronuclei with defective envelopes and impaired nuclear import^13^ **(Fig 3a)**. These envelope defects are reported to predispose micronuclei to rupture and downstream chromosome fragmentation, and to impart deficiencies in DNA replication and repair^12^. We therefore asked whether micronucleus-autophagy may selectively target such micronuclei.

**Figure 3:**
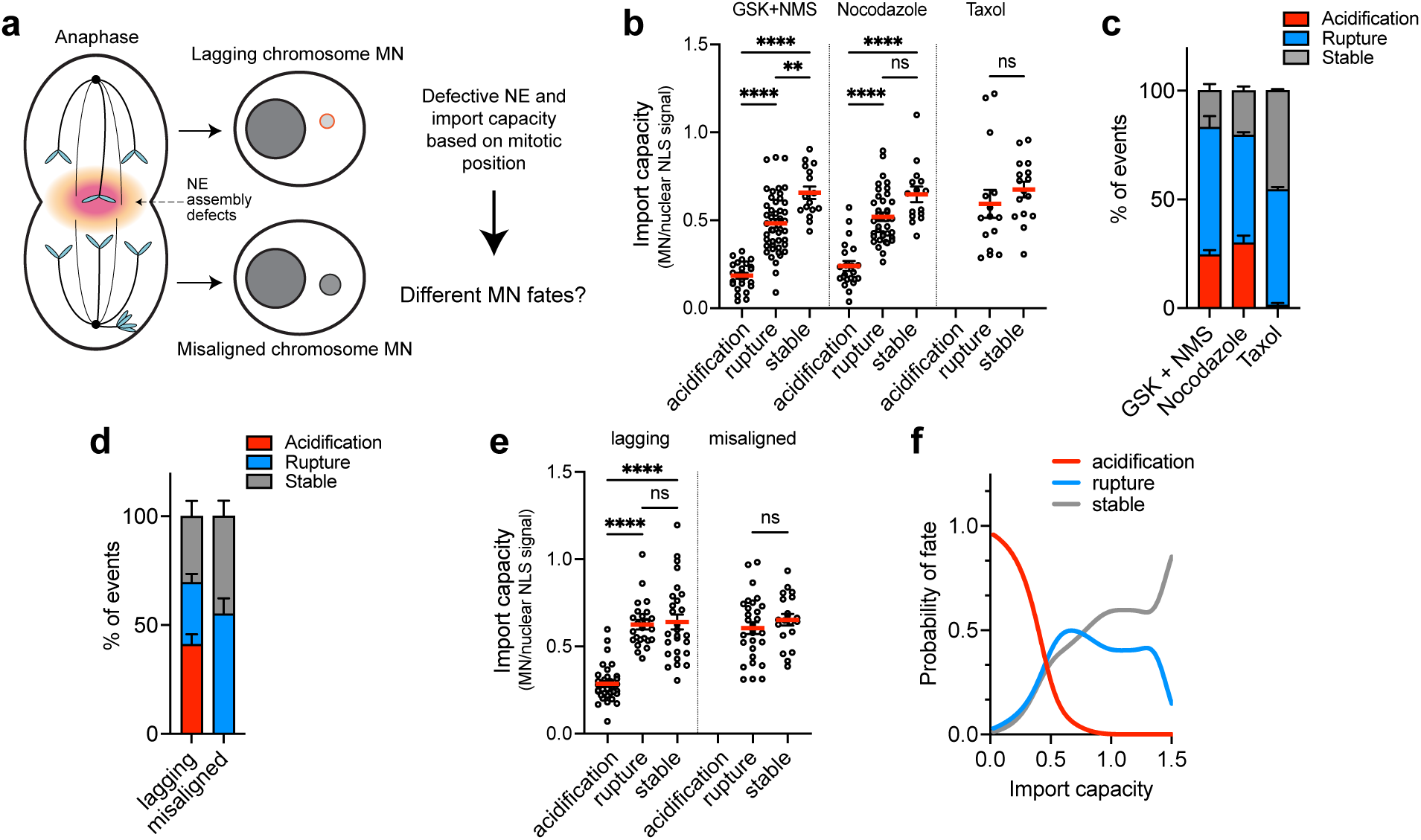
Mitotic exit dynamics predict micronucleus fate. a, Cartoon of micronuclei with nuclear envelope defects based on missegregation position, with potential impact on fates. b, Import capacity (defined as the ratio between HaloTag-NLS signal in the MN and in the primary nucleus) of MN that undergo different fates with different induction methods (n between 16 and 57 MN from n=2 experiments). c, Quantification of different MN fates with different induction methods (n≥100MN per experiment, 2 separate experiments). d, Quantification of fates for MN derived from clear lagging or misaligned phenotypes following MN induction with CENP-E and MPS1 (GSK+NMS) inhibitor treatment (n≥50MN per experiment, 4 separate experiments). e, Import capacity and fates of lagging or misaligned MN (n between 24 and 35 MN, 3 separate experiments). f, Probability of different MN fates based on their import capacity (n between 66 and 93 MN, 3 separate experiments) Data are shown as mean ± sem, and were analysed using a one-way ANOVA (b,e) (ns = non-significant, ** = p ≤ 0.01, **** = p ≤ 0.0001)

To test this, we examined the relationship between micronuclei nuclear import capacity [a measure of envelope functionality^13^] and acidification and rupture fates in hTERT RPE-1 cells expressing both a chromatin acidification and nuclear import reporter.

At variance with previous reports^13^, we observe only a modest (and in some cases non-significant) association between micronucleus import capacity and rupture frequency **(Fig 3b)**. Instead, we find that micronuclei with poor nuclear import are consistently and overwhelmingly targeted for acidification, regardless of induction method **(Fig 3b)**. Of note, micronuclei generated with low dose taxol display higher import capacity, potentially explaining the negligible micronucleus acidification rates observed in these cells **(Fig 3b, c)**.

Micronuclei which form from chromosomes positioned outside of the anaphase spindle are reported to be unaffected by nuclear envelope defects **(Fig 3a)**^13^, pointing to a relationship between chromosome missegregation phenotype and micronucleus fate. To assess this possible relationship, we used a combination treatment of CENP-E inhibition followed by Mps1 inhibition, producing a mixed population of micronuclei derived from both lagging and misaligned chromosomes. More than 40% of micronuclei originating from lagging chromosomes undergo acidification in the subsequent cell cycle, whereas those derived from misaligned chromosomes are never affected **(Fig 3d)**. By comparison, rupture events are common across both phenotypes, with particularly high rates (>50%) in micronuclei that have originated from misaligned chromosomes **(Fig 3d)**. In line with this, micronuclei derived from misaligned chromosomes have much higher levels of NLS import capacity **(Fig 3e, Fig s3a)**^13^.

Consistent with the nuclear envelope assembly defects of persistent laggards^13,37^, the acidification-fated micronuclei population is notably smaller, has lower levels of lamin B1, and shows elevated levels of emerin upon formation **(Fig S3b,c)**^13^. These micronuclei also form envelopes with moderately elevated levels of BAF at mitotic exit despite intact nuclear compartmentalisation **(Fig 2a,S3c)**, suggesting that initial BAF overrecruitment may be a feature of defective nuclear envelope assembly rather than a marker of membrane rupture **(Fig S3d)**.

Pooling our data on import capacity and micronucleus fates across experiments, we performed non-parametric Kernel density estimate (KDE) classification to determine the probability of each micronucleus fate as a function of micronucleus import capacity. We find that poor import capacity is highly predictive of micronuclei acidification, while milder import defects are associated with envelope rupture or stability **(Fig 3f).** We conclude that micronucleus-autophagy is a selective process that specifically targets micronuclei harbouring nuclear envelope defects. Consequently, the missegregation dynamics that give rise to micronuclei, and the resulting degree of nuclear envelope functionality, predict micronuclear fate outcomes.

### Chromatin-nuclear envelope untethering triggers micronucleus targeting

To investigate the mechanistic basis for the selective targeting of micronuclei with nuclear envelope (NE) defects, we next examined the localisation and dynamics of aberrantly recruited NE components in live micronuclei. We focused on three proteins, each of which is recruited to acidification-fated micronuclei at abnormal levels^13^ **(Fig S3c)**: emerin, a LEM-domain protein embedded in the inner nuclear membrane (INM); lamin B1, a component of the nuclear lamina; and BAF (Barrier-to-autointegration factor), a chromatin-binding protein that anchors chromatin to the NE. Together these markers allowed us to assess the integrity of the membrane, lamina, and chromatin-NE interface in micronuclei with and without nuclear envelope assembly defects, and according to fate.

In stable micronuclei, Emerin, Lamin B1, and BAF all maintain uniform localisation at the NE **(Fig 4b-c, S4a,c)**. Rupture-fated micronuclei (with rare exceptions) maintain the same localisation patterns up to the point of rupture **(Fig 4b-c**, **Fig 2a, S2a-c)**, upon which they display sudden, massive recruitment of BAF **(Fig 2a)**, and frequent infiltration of Emerin into the micronucleus interior **(Fig S4b**). In contrast, most acidification-fated micronuclei exhibit a pattern of progressive, sudden, or oscillating dissociation of BAF from the NE **(Fig 4a,b, S4d)**, culminating in diffuse (micro)nucleoplasm localisation shortly before autophagosomal capture **(Fig 4a-b, s4c,d)**. Lamin B1 appears to share the dissociation phenotype and dynamics of BAF **(Fig 4c, S4a,c)**, while Emerin remains stably localised to the nuclear envelope and indistinguishable from that of stable micronuclei until GFP signal quenching upon acidification^28^ **(Fig S4b)**. A sustained BAF/Lamin B1 dissociation state (defined as 3 consecutive frames of <0.8 cortical/inner micronucleus fluorescence signal), is strongly associated with acidification fate, with around 70-80% of micronuclei developing this phenotype prior to acidification **(Fig S4c)**. It is also highly predictive, with >80% of micronuclei displaying sustained BAF/Lamin B1 dissociation subsequently undergoing acidification **(Fig 4b,c)**. Sustained BAF dissociation also coincides with sudden recruitment of autophagosomal membranes to affected micronuclei, suggesting that BAF/Lamin B1 dissociation enables autophagic recognition **(Fig 4d)**.

**Figure 4:**
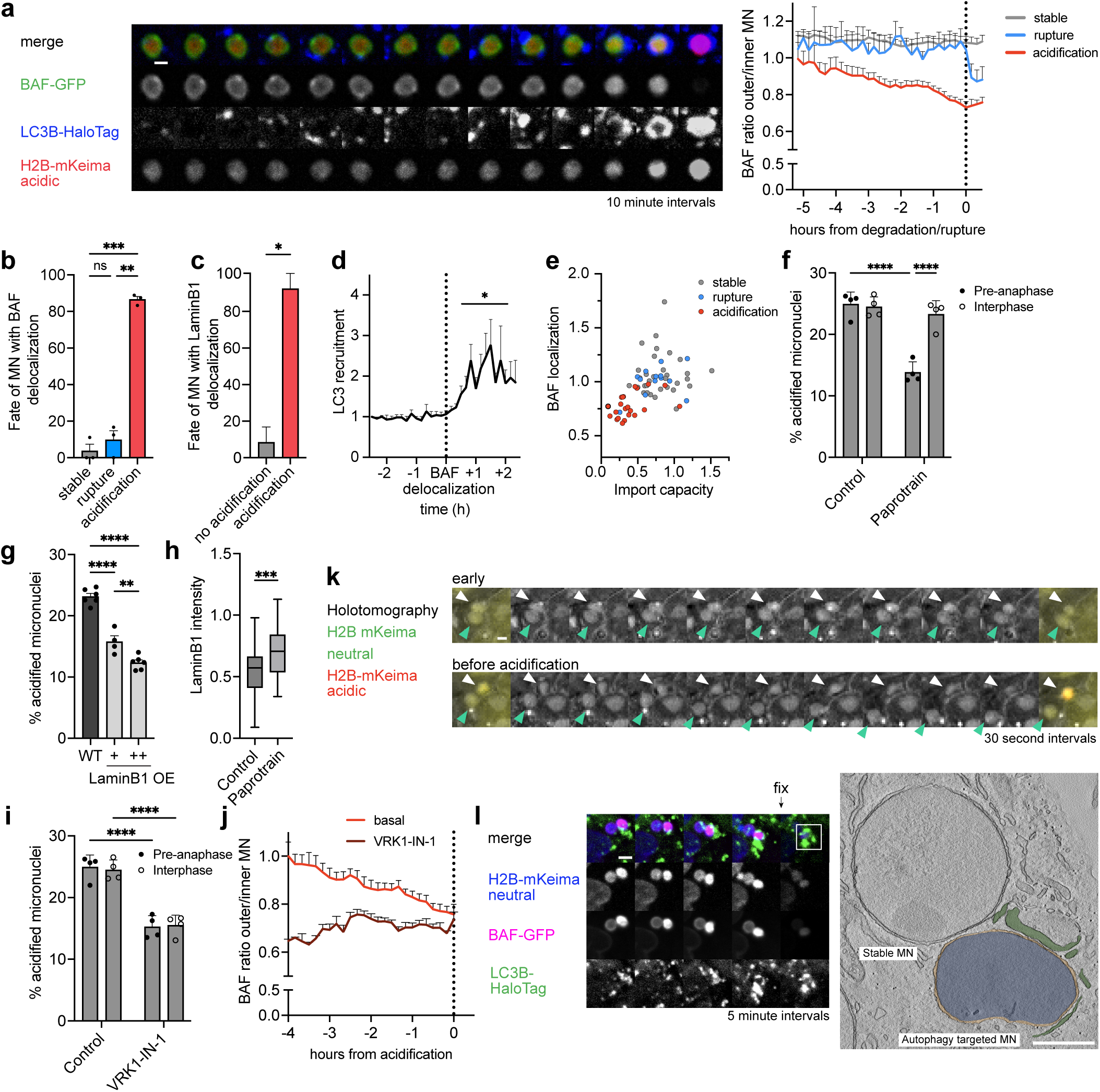
Structural defects seeded at mitotic exit trigger micronucleus targeting. a, Montage showing BAF dissociation from the MN membrane before acidification and timecourse of BAF (cortical/inner) signal for MN populations with different fates (scale bar = 1μm). b, Fate of MN with sustained (≥ 3 consecutive frames) BAF dissociation (n≥50MN per experiment, 3 separate experiments). c, Fate of MN with sustained (≥ 3 consecutive frames) Lamin B1 dissociation (n≥50MN per experiment, 3 separate experiments). d, LC3 recruitment on MN after sustained BAF dissociation (n=22 MN from 2 individual experiments). e, Relation between initial import capacity and final BAF localization based on MN fate. f, Acidification rate of MN in cells treated with the MKLP2 inhibitor, paprotrain, from nocodazole washout (pre-anaphase) or in the next interphase (n≥100MN per experiment, 4 separate experiments). g, Acidification rate of MN from cells with or without Lamin B1 overexpression(n≥100MN per experiment, 4 separate experiments). h, Lamin B1 levels on lagging derived MN in cells treated with or without paprotrain from nocodazole washout (n=50 individual MN, 2 separate experiments). i, Acidification rate of MN in cells treated with the VRK1 inhibitor, VRK1-IN-1, from nocodazole washout (pre-anaphase) or in the next interphase (n≥100MN per experiment, 4 separate experiments). j, Timecourse of BAF localization in acidified MN in basal conditions or following VRK1-IN-1 treatment from nocodazole washout (n between 14 and 21 MN from n=2 experiments). k, Holotomography montage of adjacent MN (white arrow=acidification-fated; green arrow=stable) several hours (top) and shortly (bottom) before acidification, showing rapid changes in the morphology of the acidification-fated MN shortly before acidification (scale bar = 1μm). l, CLEM experiment showing deformed but intact NE of autophagy targeted MN (green = LC3 positive autophagosomal membrane; orange = intact NE) (scale bar = 2μm for immunofluorescence image and scale bar = 500nm for EM image). Data are shown as mean ± sem, and were analysed using a one-way ANOVA (b, g), unpaired t-test (c, h), one-sample t-test (d), two-way ANOVA (f, i) (ns = non-significant, * = p ≤ 0.05, ** = p ≤ 0.01, *** = p ≤ 0.001, **** = p ≤ 0.0001)

Micronuclei which undergo BAF dissociation have almost universally established defective nuclear envelopes at mitotic exit **(Fig 4e)**, indicating that defects seeded at mitotic exit may be responsible for subsequent BAF/Lamin B1 dissociation and micronucleus targeting. In line with this, pharmacological inhibition of MKLP2 with Paprotrain, which is reported to partially rescue NE assembly defects on lagging chromosomes^38^, reduces acidification rates by 50% **(Fig 4f).** Importantly, this effect is only observed if Paprotrain is added *before* mitotic exit, underlining the mitotic origin of defects leading to BAF/Lamin B1 dissociation and micronucleus targeting **(Fig 4f)**.

Because both BAF and Lamin B1 have important roles mediating chromatin-NE tethering^39–44^ we hypothesized that their abrupt dissociation from the NE may reflect failure of these attachments. If so, interventions that strengthen chromatin-NE tethering would be expected to reduce micronuclei acidification rates.

Acidification-fated micronuclei display reduced Lamin B1 levels **(Fig S3c)**, potentially weakening their chromatin-NE tethering^44^. To explore the possibility that increasing Lamin B1 levels may strengthen tethering and reduce acidification rates we generated a Lamin B1-GFP expressing cell line and split it into high and low expressing populations **(Fig S4e)**. Micronuclei acidification rates are substantially reduced in both populations compared to the parental line, and display a clear dose response **(Fig 4g)**. Paprotrain treatment also largely rescues Lamin B1 levels on lagging chromosomes **(Fig 4h)**, potentially accounting for its suppressive effect on acidification rates.

BAF is known to mediate chromatin-NE attachments through direct binding of both chromatin and INM-embedded LEM domain proteins^39–42^. These interactions are regulated by the phosphorylation activity of the kinase VRK1, which drives global BAF dissociation during mitosis^45,46^, and promotes BAF turnover at the nuclear envelope during interphase^45^. We reasoned that suppressing VRK1 activity may stabilize chromatin-NE attachments mediated by BAF, and reduce dissociation and acidification rates. To assess this, we treated hTERT RPE-1 cells released from nocodazole arrest with a selective inhibitor of VRK1 kinase activity (VRK-IN-1) ^47^. VRK-IN-1 addition reduces acidification rates by around 50% **(Fig 4i)**. This effect is consistent whether initiated before or after nuclear envelope reformation (NER) **(Fig 4i)**, implicating interphase VRK1 activity in BAF dissociation. A comparison of initial (shortly after NER) vs final (before acidification) BAF dissociation scores of individual acidification-fated micronuclei confirms progressive deterioration of BAF NE-localisation for a large portion of the acidification-fated population **(Fig 4j, s4f)**. VRK1i largely rescues this deterioration, with the remaining population of acidification-fated micronuclei displaying critically NE-dissociated BAF upon formation **(Fig 4j, s4f)**.

Chromatin-NE tethering contributes substantially to nuclear stiffness and viscosity^48^. If the observed BAF/Lamin B1 dissociation phenotype reflects failure of these attachments, affected micronuclei would be expected to display an increase in NE fluidity and deformation susceptibility. In support of this interpretation, rapid-interval holotomographic/fluorescence imaging of 2 adjacent micronuclei with well resolved membranes illustrated a dramatic increase in the NE membrane fluidity and mobility of the acidification-fated micronuclei shortly before acidification **(Fig 4k, Supplementary video 1)**. Furthermore, CLEM analysis of adjacent micronuclei fixed after one had incurred BAF dissociation and LC3 targeting (but not acidification), also illustrated a deformed but intact NE targeted by large autophagosomal membranes **(Fig 4l)**.

Taken together, our data support a model in which defects seeded at mitotic exit lead to progressive loss of chromatin–NE tethering, producing a mechanically altered state that is selectively recognised by the autophagic machinery.

### Clearance of micronucleus entrapped chromosomes

To examine the genomic consequences of micronucleus-autophagy, we labelled cells with a live-cell DNA dye and monitored acidification events. Micronucleus acidification is marked by a sudden increase in DNA dye fluorescence signal localized on the micronucleus, followed by progressive signal loss over a period of approximately 2 hours, after which the micronuclear DNA dye becomes undetectable **(Fig 5a, S5a)**. DNA staining of cells fixed after a live-imaging-verified acidification event shows a similar decay in micronuclear DNA signal over 1-2 hours, but without the initial fluorescence increase (which we presume to be an artifact of pH-dependent dye behaviour in live cells). A similar pattern is observed when DNA content is assessed by immunofluorescence of incorporated EdU **(Fig 5b).** Furthermore, TUNEL staining of live-imaging–verified acidification events reveals transient detection of DNA strand breaks, coinciding with the acute loss of Hoechst DNA dye signal. **(Fig. 5c)**. Collectively, these results suggest rapid degradation of entrapped chromosomal material within lysosomes.

**Figure 5:**
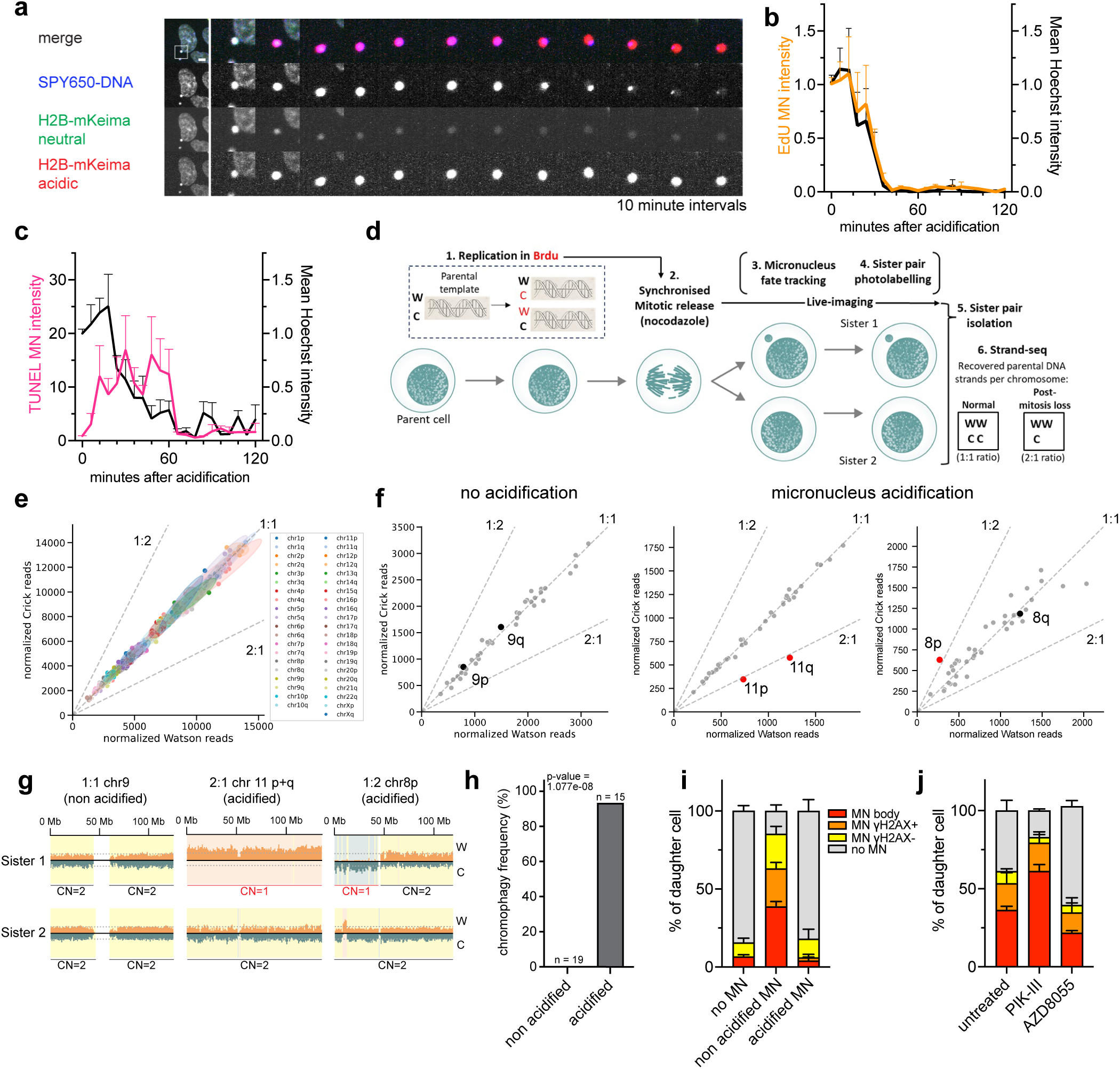
Clearance of micronuclei entrapped chromosomes by chromophagy suppresses chromosomal instability. **a,** Montage showing MN acidification followed by loss of DNA dye signal (scale bar = 5μm). **b,** Timecourse of EdU and Hoechst signal intensity after MN acidification (n=83 acidified MN and n=33 non acidified MN, 3 separate experiments). **c,** Timecourse of Hoechst and TUNEL signal intensity after MN acidification showing transient detection of broken DNA ends (n=99 acidified MN and n=35 non acidified MN, 3 separate experiments). **d,** Schematic of fate-resolved sister-cell strand-seq. Brdu incubation results in each sister chromosome receiving 1 non-Brdu incorporated (parental) DNA strand, either W or C. Sister cells with known MN fates (determined by live-imaging) are photolabelled and isolated by FACS (methods). Following Brdu strand degradation and single-cell sequencing, unbalanced W:C ratios between sister chromosomes are only explainable by post-mitotic loss of a sister chromosome. **e,** W:C chromosome arm ratio plot from 8 aggregated previously 6 sequenced (untreated) sister cell pairs with normal nuclei, showing overall balanced ratios. **f,** representative W:C chromosome arm ratio plots for live-imaging–validated sister cell pairs containing either non-acidified (left) or acidified micronuclei (middle, right). Highlighted chromosomes, shown in black or red according to whether they correspond to scenario 1 or scenario 2, are those displayed in panel g. **g,** W:C strand reads mapped to highlighted chromosomes from (5f), non acidified (left), acidified (middle, whole chromosome loss), acidified (right, chromosome arm loss). **h,** sister cell pairs with de novo chromosome or chromosome arm level loss by live-imaging verified MN fate. **i,** MN and MN bodies in daughter cells by parental cell MN status and fate. **j,** MN and MN bodies in the daughters of MN +ve parental cells following pharmacological autophagy inhibition (PIK-III) or stimulation (AZD8055).

Multiple prior studies have linked micronuclei to accelerated karyotypic evolution in cancer, particularly via chromosomal alteration (CA) processes characterized by extensive DNA fragmentation and rearrangement^5–7,33^. To evaluate the impact of micronucleus-autophagy on CA formation, we combined live-cell micronucleus fate tracking with DACT-1–based photoactivation (**Methods**) to isolate sister cell pairs containing micronuclei with known fates. We profiled recovered cells using template strand sequencing (Strand-seq)^49^, a haplotype-resolved single-cell genome sequencing approach that allows the reconstruction of reciprocal *de novo* CA trajectories across sister cell pairs based on parental template strand information^6,50^ (**Fig 5d**). We confirmed sister cell relationships from the template strand data^6^ (**Methods**), allowing us to assess the fate of all chromosomes even in the context of frequent mis-segregation events between sister pairs. We identified 15 sister pairs which had exhibited a live-imaging verified micronucleus acidification event, and 19 pairs in which acidification did not occur. As expected, we observe frequent reciprocal *de novo* whole chromosome and chromosome arm level aneuploidies between sister pairs, reflecting Nocodazole induced missegregation events^6,51^. We therefore split chromosomes into p and q arms for further downstream analysis.

In analysing these single cell data, we considered two distinct scenarios for entrapped chromosomes within acidified micronuclei. Under *scenario 1*, chromosomes confined within micronuclei remain intact. To generate baseline data representative of this scenario, we analyzed single cell data from 8 untreated sister cell pairs with normal nuclei, which are presumed to retain their original karyotype without *de novo* CAs^6^ (**Methods**). Quantification of Watson (W) and Crick (C) parental-template strands across each pair shows a balanced distribution for every chromosome, with W:C ratios approximating 1:1 when template strands are aggregated across sisters **(Fig 5e**). This is consistent with the classical Watson–Crick model of DNA replication, in which equal division of replicated sister chromatids results in each daughter cell inheriting one parental strand (W or C) of the original template duplex^52^. This results in an equal representation of W and C strands between sister chromosomes (here used to denote chromosomes originating from sister chromatids) across a sister cell pair **(Fig 5d)**.

Under *scenario 2*, we considered the possibility that after DNA replication and mitosis, a single chromosome becomes confined to an acidified micronucleus, in which it is fully degraded. If micronucleus-autophagy triggers chromosome digestion, the predicted consequence would be a skewed W:C ratio of parental-template DNA, ∼1:2 or ∼2:1 (in the case of disomy), for only this chromosome within otherwise W:C ratio-balanced sister chromosomes. Such a skew would appear to violate the Watson–Crick model of equal template inheritance^52^, with this imbalance only explainable by post-mitotic elimination of a single sister chromosome **(Fig 5d)**.

Analysis of 19 sister cell pairs in which micronucleation occurred *without* acidification show uniform W:C ratios near 1 for almost all chromosomes **(Fig. 5f,g, S5b)**. While some chromosome arms have apparent outlier W:C ratios (evident in 6 of the pairs), inspection of mapped reads clearly reveals these to be missegregated but intact (with the skew resulting from normalisation between sisters with different read depth) **(Fig. S5b)**. Thus 19 of 19 pairs correspond to scenario 1.

By comparison, among 13 out of 15 sister pairs which incurred a live-imaging verified micronucleus acidification event we observe at least one chromosome or chromosome arm clearly corresponding to scenario 2 (1:2 or 2:1 W:C ratio). As a complementary analysis approach, we also performed automated copy-number calling across sister cell pairs using strandtools (**Methods**). This corroborated our W:C ratio analysis and highlighted an additional non-reciprocal chromosome loss among the acidified pairs (which was likely not detected based on W:C ratio due to disparate read depths between sister cells). Thus, in total, 14 of 15 cell pairs which incurred MN acidification and 0 of 19 pairs with non-acidified MN contain a whole chromosome or chromosome arm loss which is non-reciprocal and thus corresponds to scenario 2 (*P*=1.08e-08, two-tailed Fisher’s exact test) **(Fig. 5f-h, S5c).** Several of these pairs also harbor multiple independent events affecting different chromosomes, for a total of 11 whole-chromosome and 6 chromosome-arm losses, compared with 0 such events observed across 19 sister-cell pairs lacking detectable acidification **(Fig. S5b-d)**. Non-acidified sister cell pairs contain frequent whole chromosome and chromosome arm level copy-number changes, but these are essentially always reciprocal between sister cell pairs, with gains in one sister mirrored by loss in the other **(Fig S5e)**. In contrast, sister cell pairs which underwent micronucleus acidification show significantly more frequent non-reciprocal events (*P*=1.87e-05, two-tailed Fisher’s exact test), and a loss:gain ratio near 2:1 for both whole chromosome and chromosome arm level copy number changes **(Fig S5e)**.These results indicate that degradation of DNA derived from one parental template occurs specifically in sister pairs displaying micronuclear acidification. Therefore, DNA within acidified micronuclei is efficiently eliminated, resulting in loss of large-scale chromosomal material, including entire chromosomes.

There are immediate implications arising from these data. These findings suggest that distinct micronuclear fates differentially influence the spectrum of emerging CA classes. In particular, micronucleus-autophagy, unlike other fates, results in the effective degradation of the entrapped DNA, causing *de novo* loss of genomic material that subsequently is no longer detectable within a sister cell pair. By facilitating the cellular clearance of chromosomes, this process may also limit the transmission of missegregated chromosomal material to daughter cells. In light of these findings, we propose the term chromophagy (Greek: *chromo*, chromosome; *phagy*, eating) for this previously uncharacterized process.

### Chromophagy arrests the intergenerational transmission of micronuclei

CAs originating from micronuclei develop through the repeated transmission, fragmentation and progressive structural rearrangement of chromosomes across successive cell cycles^5,6,15^. To investigate the consequences of chromophagy on this process, we tracked fate-matched micronucleated cells through a subsequent cell division and stained for γH2AX in daughter cells. We find that 50% of daughter cells descending from micronucleated cells that did not undergo chromophagy also contain micronuclei, and a majority of these are heavily stained for γH2AX, indicating that they contain fragmented chromosomal material that appears to persist and propagate over successive cell divisions **(Fig 5i, S5f)**. A further 40% of daughter cells display large γH2AX territories within the primary nucleus, previously termed MN bodies^53^, characteristic of reincorporated micronucleus-derived chromosome fragments **(Fig 5i, S5f)**. In contrast, micronucleated cells subject to chromophagy produce daughter cells with minimal levels of micronuclei and MN bodies, similar to background levels **(Fig 5i)**. We conclude that chromophagy arrests the intergenerational transmission of micronuclei-derived chromosomal material.

In line with these findings, intergenerational tracking of cells treated with an autophagy inhibitor (PIK-III) or autophagy stimulator (AZD8055) shows that stimulation of chromophagy reduces the abundance of MN bodies, while inhibition of chromophagy leads to their accumulation **(Fig 5j)**. These treatments had no impact on cells without MN, confirming that the observed effect is due to MN degradation **(Fig S5g)**.

In summary, chromophagy clears cells of micronuclei-entrapped chromosomal material and arrests the intergenerational transmission of micronuclei derived chromosome fragments.

## Discussion

Micronuclei-entrapped chromosomes present a unique liability to genomic integrity due to their vulnerability to iterative CA formation over multiple cell cycles^2,16,17^. Although cells possess numerous checkpoints and safeguards that promote faithful chromosome segregation and limit micronucleation and aneuploidy, the extent to which they can mitigate the genome-destabilizing consequences of micronuclei once formed has remained unclear. In this work we describe chromophagy, a selective autophagy pathway that targets whole micronuclei for degradation. Our data indicate that chromophagy efficiently removes chromosomal material entrapped within micronuclei from the genome. Lysosomal nucleases, such as lysosomal DNase IIα^54,55^, might mediate chromosomal degradation during chromophagy. Through this process, chromophagy counteracts micronuclear rupture, prevents chromosomal fragmentation and complex rearrangements, and restricts the transmission of micronucleus-derived chromosomal fragments to daughter cells. This limits the downstream genomic consequences of micronucleation, and as such could suppress rapid karyotype evolution^2,16,17^ involving mutational processes such as breakage-fusion-bridge cycles and chromothripsis.

An important feature of chromophagy is the strong selectivity for micronuclei exhibiting defects in nuclear envelope assembly and function, characteristic of micronuclei formed from persistently lagging chromosomes. These defects have previously been linked to micronucleus rupture and downstream chromosome fragmentation, and were therefore considered a deleterious byproduct of chromosome missegregation^13^. Instead, our data indicate that these defects seed weaknesses in chromatin-NE attachments, promoting interphase chromatin-NE dissociation, and marking micronuclei for chromophagy. Viewed in this way, the assembly of defective nuclear envelopes on missegregated chromosomes could constitute an additional segregation error correction mechanism, analogous to, and possibly instantiated by, the anaphase surveillance activity of midzone localised Aurora B^38,56,57^.

Defining the “ eat-me” signals that mark these micronuclei for autophagic elimination will be essential for understanding how selective recognition is achieved. Our findings suggest that loss of chromatin-NE attachments, dependent on both defects seeded at mitotic exit and VRK1 activity during interphase, generates a distinctive mechanical configuration on the surface of targeted micronuclei. This transition could create a permissive interface for the recruitment of autophagic factors; however, the precise molecular determinants of this recognition remain to be defined. One possibility is that mechanotransductive changes at the micronuclear surface expose normally shielded protein or lipid moieties that function as damage-associated signals. Consistent with this view, our data show that chromophagy requires the canonical autophagy core machinery and membrane-associated recruiters of cargo adaptors from the ATG8 protein family, indicating that micronucleus recognition is mediated through established pathways of selective autophagy. Clarifying the molecular nature of these signals will therefore be crucial for understanding how aberrant nuclear structures are identified and eliminated by chromophagy before they can propagate genome instability.

### Safeguarding genomic integrity through arresting CIN

Our finding that chromophagy removes micronucleus-entrapped chromosomes suggests that this process might contribute to genome surveillance with potential tumor-suppressive consequences. Our experiments were largely pursued in p53-proficient hTERT RPE-1 cells, a diploid, non-transformed cell model. In such a setting, where checkpoint control and autophagic capacity are essentially intact, chromophagy is likely to act within a broader network of genome-protective mechanisms. Indeed, autophagy is presumed to suppress tumor initiation in early stages of cancer development by sustaining genomic integrity^58^, including through apoptosis during replicative stress, centrosome quality control, oncogene-induced senescence, and removal of DNA lesions to promote DNA repair^21,59–62^. In comparison to these previously described mechanisms, chromophagy may specifically mitigate detrimental genomic consequences of micronucleation by constraining the emergence of chromothripsis and other CA processes^2^. As such, chromophagy may act as a barrier to karyotype destabilisation linked to mitotic errors.

It is also possible that chromophagy contributes to suppressing tumor development by limiting the non-genetic effects of micronucleation. Micronuclei entrapped chromosomes undergo epigenetic alterations accompanied by heritable transcriptional changes, and these modifications can persist during clonal expansion, which has been proposed to promote the formation of transcriptional heterogeneity in cancer^53^. We find that chromophagy selectively targets defective micronuclei, presumed to harbor the most substantial epigenetic abnormalities^53^. By preventing the transmission of epigenetically altered chromosomes, chromophagy may limit the formation of transcriptionally diverse cancer cell subpopulations that contribute to tumour progression, metastasis and drug resistance^63–65^.

There is also the possibility that, in certain settings, chromophagy might facilitate tumour maintenance or progression. Autophagy has dual roles in cancer biology, acting either in a suppressive or oncogenic manner depending on cellular context^66^. A prevailing hypothesis posits that genomic instability drives tumour development only below a critical limit, since excessive CIN can result in deleterious karyotypes reducing cellular fitness^3,67–69^. Our results from U2OS and HeLa cells indicate that chromophagy is operative in established cancer cells. By constraining the iterative accumulation of complex CAs, chromophagy may buffer CIN at levels supporting sustained cancerous growth.

### Chromophagy results in a net mutational bias towards chromosomal loss

While chromophagy suppresses micronucleation-mediated CIN, it simultaneously leads to the effective loss of genomic material. Pan-cancer analyses have consistently identified an intriguing imbalance in cancer genomes, with a predominance of whole-chromosome losses over gains^6,24,25^. While selective forces acting during cancer evolution might preferentially retain chromosome losses, data from a study coupling imaging and single-cell sequencing in different human cell lines showed that chromosomal losses arise more frequently *de novo* than gains in micronucleated cells^6^. Since chromophagy promotes degradation of DNA, including of entire chromosomes, it offers a potential mechanistic explanation for this mutational asymmetry.

Indeed, chromophagy also likely plays a role in shaping the aneuploidy landscape of cells experiencing CIN. The impact on aneuploidy will depend on whether targeted micronuclei reside within the correct or incorrect sister with respect to the maintenance of ploidy. In the case of a micronucleus-entrapped chromosome residing in the incorrect sister (giving rise to a trisomic state), chromophagy may serve to eliminate the additional chromosomal copy and thus counteract trisomy formation. This would effectively halve the aneuploidy rate between this sister pair. On the other hand, eliminating a micronucleus-entrapped chromosome residing in the correct sister would come at the cost of introducing a monosomy. Thus while chromophagy will always have the effect of suppressing ongoing micronucleus-mediated CIN, its impact on aneuploidy rates depends on context. Importantly, in all cases the net impact across sister cell pairs will be the predominance of chromosomal losses over gains. Whether chromophagy rates vary between micronuclei residing within correct or incorrect sister cells is an interesting question for future study.

### Therapeutic potential of chromophagy

From a therapeutic perspective, modulation of chromophagy might offer potential strategies to constrain rapid karyotype evolution. If chromophagy limits the iterative formation of complex chromosomal rearrangements over successive cell cycles, its pharmacological activation might reduce the generation of karyotypic diversity fueling clonal selection under therapeutic pressure^3^. For instance, chromophagy-enhancing interventions might be deployed as adjuvant treatments to stabilize tumour genomes during chemotherapy or targeted therapy, thereby suppressing the emergence of resistant subclones. Such an approach would not aim at direct cytotoxicity, but rather at restricting the evolutionary trajectories available to cancer cells, potentially prolonging response duration and delaying relapse.

## Acknowledgements

We thank Vanda Turcanova for cloning support and Patrick Hasenfeld for extensive support with FACS isolation of sister cells and single-cell sequencing library preparation. We are grateful to the Danish Cancer Institute and EMBL core facilities and staff for technical support, including imaging, flow cytometry, and single-cell sequencing support. We also thank members of the Korbel and Cecconi groups for helpful discussions and feedback throughout the project. Funding for this work came from the European Research Council (ERC Advanced grant (SEE-MAGIC) grant no. 101098056) to J.O.K.; from a Health and Life Science Alliance postdoctoral fellowship to N.A.W; from a Dansk Kræftforskningsfonds grant (ID 2025-65-1175) to M.M.; from Fondazione AIRC per la Ricerca sul Cancro (AIRC) under grant IG-2025-32370, the Novo Nordisk Foundation grant no. 0094651, and the Italian Ministry of University and Research (MUR) under the PRIN 2022 PNRR programme funded by the European Union – NextGenerationEU (Project code P2022LZP9T) to F.C.

## Data

Genomic datasets generated and analysed in this study are available from the European Nucleotide Archive (ENA) under accession numbers PRJEB109929 (newly generated data) and PRJEB78885 (previously published data^6^ re-analysed in the present study).

## Author contributions

N.A.W. conceived key aspects of the study and led experimental design, data acquisition, and analysis. Conceptual input on autophagy pathway regulation was provided by F.C., and conceptual input on the investigation of genome instability consequences was provided by J.O.K. M.M. performed experiments and contributed to data analysis and manuscript writing. M.R.C. and J.O.K. developed and implemented single-cell sequencing analysis methodology. V.O. performed electron microscopy experiments. L.B.F. contributed to supervision and scientific discussion. F.C. and J.O.K. provided senior supervision, guidance and resources. N.A.W., J.O.K. and F.C. wrote the manuscript.

## Material and methods

### Statistical analysis

Data are generally shown as mean ± sem and significance level is indicated as follows: ns = non-significant, * = p ≤ 0.05, ** = p ≤ 0.01, *** = p ≤ 0.001, **** = p ≤ 0.0001.

Statistical tests (performed in GraphPad Prism 10.4, unless otherwise indicated) are indicated in the figure legends and were selected based on data distribution and experimental design. Parametric tests were used for normally distributed data, while non-parametric tests were applied otherwise. Comparisons between a single group and a reference mean were performed with a one-sample t-test, comparisons between 2 groups were performed using a t-test, while one-way ANOVA was used for comparisons among three or more groups. Two-way ANOVA with post hoc multiple comparison tests was applied when two independent variables were considered.

Kernel density estimation (KDE) analysis was performed in R (version 4.3.2). Data were arranged to correlate a numerical variable (Import capacity) and a categorical variable (MN fate). Probability densities were estimated using Gaussian KDE; the analysis’s bandwidth was selected based on visual optimization to balance smoothing and feature retention. Densities were normalized across fate classes to represent mutually exclusive probabilities. Multinomial logistic regression was used to model the probability of each fate as a smooth function of the measured variable.

### Cell culture

hTERT RPE-1, U2OS, HeLa and HEK293T cells were purchased from the American Type Culture Collection (ATCC) and cultured in Dulbecco’s Modified Eagle’s Medium (DMEM) GlutaMAX (Gibco) supplemented with 10 % fetal bovine serum (FBS) (Gibco) and 1% penicillin/streptomycin.

All cell lines were grown at 37 °C in a humidified incubator with 5% CO2.

Stably integrated construct expression was induced with 48h treatment with doxycycline (500ug/mL, Sigma-Aldrich D9891)

### Plasmids

pFUW-HaloTag-NLS was generated using the Gibson Assembly mix (NEB) to substitute mCherry from pFUW-TetO-mCherry-cNLS (Addgene Plasmid #138531) with HaloTag coming from pMRX-No-HaloTag7-LC3 (Addgene Plasmid #184900). pLVX-TetOne-H2B-mKeima[MM2] was generated using the Gibson Assembly mix (NEB) to substitute TOMM20 from pLVX-TOMM20-mKeima with H2B coming from H2B-EGFP-IRES-ER-dKeima (Addgene Plasmid #69550). EGFP-BAF was a gift from Daniel Gerlich (Addgene plasmid # 101772); pLVX-EF1a-EGFP-Emerin-IRES-Hygromycin was a gift from David Andrews (Addgene plasmid # 134864); pQCXIP-GFP-LmnB1 was a gift from John Maciejowski (Addgene plasmid # 164249); pMRX-No-HaloTag7-LC3 was a gift from Noboru Mizushima (Addgene plasmid # 184900). Plasmid containing H2B-GFP-RFP and pLVX-GFP-LC3B were obtained from Vectorbuilder.

### Lentivirus generation and stable cell line generation

Lentiviral vectors were produced by transfecting human embryonic kidney (HEK) 293T cells with MISSION® Lentiviral Packaging Mix (Sigma-Aldrich, SHP001) and the plasmid of interest using Fugene (Promega, E2311) according to the manufacturer’s protocol. Lentiviral vectors were collected after filtration of the media with 0.45 μm filter and mixed with 8 μg/ml polybrene (Sigma-Aldrich, #107689) to infect target cells. Transduced cells were selected by antibiotic selection or FACS sorting.

### ATG7 KO generation

hTERT RPE-1 H2B-mKeima were transfected using lipofectamine 3000 according to the manufacturer protocol and plasmids designed for CRISP-Cas9 mediated knock-out of ATG7 (SantaCruz, sc-400997-KO-2). Single cells were sorted into 96 well-plates, grown into clones, and screened for expression of ATG7 by western blotting.

### Western blotting

Cell lysates were prepared with whole-cell lysis buffer (50 mM Tris-HCl pH 6.8, 10 % Glycerol, 2% SDS) for 10 min at 95°C at 1200 rpm agitation. Protein extracts were quantified using the DC protein assay (Bio-Rad 500-0113 and 500-0114) and denatured in Sample Buffer. Proteins were subjected to SDS-PAGE on 4 to 15% Criterion TGX Stain-Free Protein Gels (Bio-Rad) and transferred to polyvinylidene fluoride (PVDF) membranes using semidry transfer with the Trans-Blot Turbo Transfer System (Bio-Rad).

The membranes were blocked using 5% dry milk in PBS-Tween-20 0.1% and incubated with the primary antibodies in blocking solution overnight at 4°C, followed by incubation in secondary horseradish-peroxidase (HRP)-conjugated antibodies (Bio-Rad) (1:3000) for 1 hour at RT. Secondary antibody detection was performed using Amersham ECL Prime (Cytiva, RPN2232) or Immobilion ECL Ultra Western HRP Substrate (Merck Millipore, IBULS0100) and the signal was acquired by ChemiDoc MP System (Bio-Rad).

### Antibodies

The following antibodies were used. ATG16L1 (Cell Signaling, 8089S, WB 1:1000), ATG7 (Cell Signaling, 8558S, WB 1:1000), LC3B (Novus Biologicals, NB100-2220, WB 1:1000), GABARAP (Cell Signaling, 13733S, WB 1:1000); yH2AX (Merck, 05-636, IF 1:1000), Lamin B1 (abcam, ab16048, IF 1:750)

### RNA extraction, retro transcription and real-time PCR

RNA was extracted using the NucleoSpin RNA kit (Macherey-Nagel, 740955) following the manufacturer’s protocol. cDNA was produced using M-MLV reverse transcriptase (Promega, M5313). qPCR was performed on a ViiA 7 machine (ThermoFisher) using PowerUp SYBR Green Master Mix (ThermoFisher, A25742) following the manufacturer’s protocol.

Employed primers are as follows:

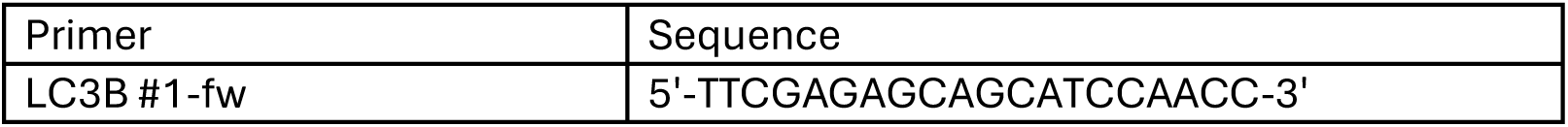
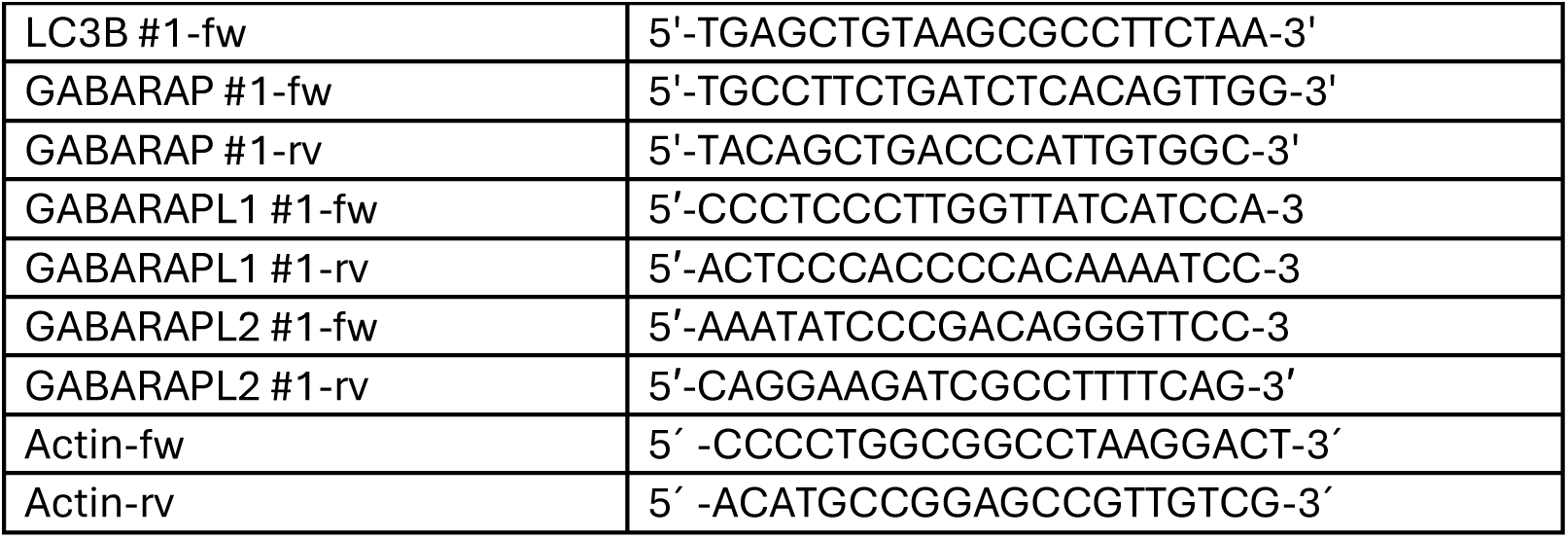

### siRNA mediated silencing

siRNA transfections were carried out using Lipofectamine RNAiMAX (Invitrogen, 13778150) adapting the manufacturer’s protocol to work with a T25 culture flask. Appropriate siRNA negative controls were used.

Employed siRNA are as follows:

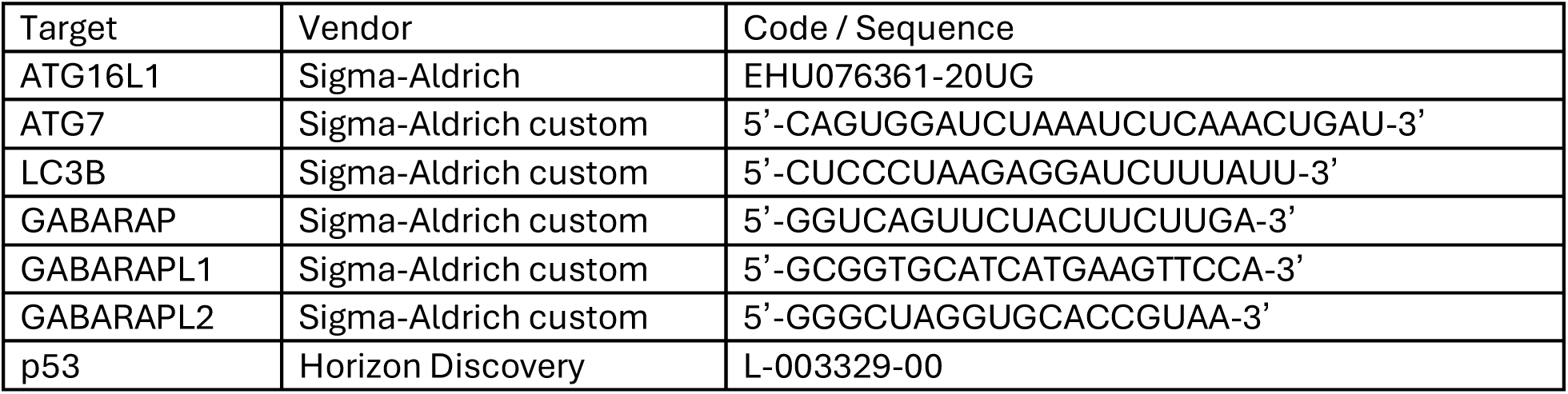

### Micronuclei induction

Most experiments used a nocodazole block and release procedure to generate micronuclei. Cells were treated with 100 ng/ml nocodazole (Sigma-Aldrich, M1404-2MG) for 6 h to arrest them in mitosis, they were then washed 3 times with pre-warmed PBS and collected via mitotic shake-off to be plated in complete medium for live-imaging. To better distinguish between lagging and misaligned chromosomes, MN were induced by arresting cells for 6h at metaphase with 75 nM of the CENP-E inhibitor GSK-923295 (MedChemExpress, HY-10299), collecting them by mitotic shake-off and plating in MPS1 inhibitor NMS-P715 (1µM, Millipore Sigma, 475949) to bypass the spindle assembly checkpoint and allow mitotic progression. Paclitaxel treatment (48h 10nM, Sigma-Aldrich, 580555) was also used as an alternative MN induction method with the drug diluted in the medium during imaging.

### Drug treatments

In live-imaging experiments drugs were, unless otherwise indicated, added to the cells after anaphase completion (2 hours after mitotic shake-off) to avoid possible unrelated effects during anaphase progression and mitotic exit and left in the medium during imaging. To modulate autophagy the following drugs were used: Chloroquine (40µM, Sigma-Aldrich, C6628), PIK-III (5µM, Cayman Chemical Company, 17002), Bafilomycin A1 (200nM, Santa Cruz Biotechnology, SC-201550), AZD8055 (100nM, MedChem Express, HY-10422), Torin1 (100nM, Selleck Chemicals, S2827), EBSS (replacing the DMEM with EBSS after 2 washes with PBS). To investigate the role of VRK1 kinase activity on BAF dissociation and MN fates VRK1-IN-1 (500nM, MedChemExpress, HY-126542) was added from nocodazole washout or after mitotic exit. To investigate the impact of suppression of Aurora B midzone localisation on MN fates and Lamin B1 levels Paprotrain (10µM, MedChemExpress, HY-101298) was added from nocodazole washout or after mitotic exit.

### Dyes for live imaging

LysoTracker™Deep Red (ThermoFisher, L12492, 50nM), SPY650-DNA (Spirochrome, SC501, 50nM) and Janelia Fluor 646 HaloTag Ligand (Promega, GA1120, 20nM) were added to the cells 4 hours after nocodazole treatment and washed out during the mitotic shake-off.

### Micronucleus fate live imaging

For live imaging experiments to quantify micronucleus fates, a Zeiss CellDiscoverer 7 with 37°C incubation and 5% CO^2^ was used. After mitotic shake-off cells were plated in 96 well plates (PhenoPlate, Revvity, 6055302) using FluoroBrite DMEM (ThermoFisher, A1896701) and imaged in the heated chamber of the microscope. Widefield fluorescent tiled images were taken with the 20x air Plan-Apochromat 0.95 NA every 20-30 minutes for 16-18h. At least 100 MN per replicate were randomly selected from the first frame and fates visually scored through the time-lapse acquisition based on the following definition criteria. Rupture events were defined as MN with a permanent and sudden loss of NLS signal. Degradation events were defined as MN with a permanent and sudden switch of the mKeima signal from the neutral to the acidic channel. Only cells containing ≤ 2 micronuclei were considered in all experiments.

### High-resolution fluorescence live imaging

For high-resolution live imaging experiments in which the main readout was the localization of factors on micronuclei, a Nikon Eclipse Ti equipped with Crest X-Light V3 or a Nikon CSU-W1 Sora spinning disc microscope was used. After mitotic shake-off cells were plated in Ibidi chambers (Ibidi, 80806) using FluoroBrite DMEM (ThermoFisher, A1896701) and imaged in the heated chamber of the microscope (37°C incubation and 5% CO^2^). Images were acquired using a CFI Plan Apochromat VC 60XC 1.2 NA WI objective (Nikon Eclipse), or a SR P-Apochromat IR AC 60x WI/ NA 1.27 (Nikon CSU-W1 SORA) objective every 5-10 minutes for 12h, with 0.5µm z-intervals.

### Analysis of high-resolution fluorescence live imaging

Z-stack images were converted to max projections in Fiji and individual MN tracks were created using the TrackMate plugin^70^.

A custom pipeline was used to analyse the localization of the different proteins. A first MN mask was created based on the signal of the POI (protein of interest) and eroded to obtain a second mask formed by only the inner part of the MN (the nucleoplasm), a third mask, comprising the cortical part of the MN (corresponding to the membrane part) was obtained by subtraction of the second mask to the first one. The average intensity of the POI was calculated for each mask after background subtraction, and the ratio between the third mask and the second one was considered.

For analysis of LC3 recruitment, the first mask was based on the H2B-mKeima channel and expanded to cover the area in the vicinity of the MN. LC3B mean intensity was calculated in the second mask after background subtraction.

### Holotomography

Holotomographic imaging was performed on a Nanolive 3D Cell Explorer Fluo **(Fig 4k, supplementary video1)**, or a Tomocube HT-X1 **(Fig 1h, Fig s2c).** hTERT RPE-1 cells expressing H2B-mKeima (and BAF-GFP for **Fig s2c)** were released from nocodazole arrest and imaged at 30s intervals on the holotomography channel and 5 minute intervals on fluorescent channels.

### CLEM

hTERT RPE-1 cells expressing H2B-mKeima, BAF-GPF, LC3-HaloTag, were released from nocodazole arrest onto ibidi chamber slides and imaged with a Nikon CSU-W1 Sora spinning disc microscope (5 minute intervals). Micronuclei were monitored for autophagic targeting, indicated by the transition to a diffuse BAF signal and LC3 targeting. Upon LC3 recruitment cells were fixed by gently adding 200 µl of fixative consisting of 6% paraformaldehyde (PFA) in 0.1 M PHEM buffer. After fixation, brightfield images and Z-stacks were acquired, and regions of interest (ROIs) were recorded. Following imaging, the initial fixative was replaced with fresh fixative composed of 2% glutaraldehyde (GA), CaCl₂, and MgCl₂ in 0.1 M sodium cacodylate buffer (pH 7.4). Samples were fixed overnight at 4 °C. After fixation, most of the fixative was removed while ensuring the samples did not dry. Fresh 0.1 M sodium cacodylate buffer was added carefully, and samples were rinsed six times for 10 min each. Post-fixation was performed in 1% osmium tetroxide with 1.5% potassium ferricyanide in 0.065 M sodium cacodylate buffer (pH 7.4) for 2 h at 4 °C in the dark. Following post-fixation, samples were rinsed thoroughly in distilled water six times for 10 min each. Dehydration was performed through a graded ethanol series of increasing concentrations. Following dehydration, samples were infiltrated with Epon 812 resin using a stepwise infusion of 25%, 50%, 75%, and 100% Epon. After the final infiltration step, most of the resin was removed from the well, and a Beem capsule filled with fresh liquid resin was positioned upside-down onto the sample. Polymerization was initiated by incubating the samples at 66 °C for 24 h. Beem capsules were detached from the Ibidi chamber wells, leaving the 180 µm Ibidi plastic layer still attached to the surface of the resin block. Polymerization was continued for an additional 48 h at 66 °C. Polymerized resin blocks were trimmed, on the sides, to the region of interest using a Leica Trim2 system. The 180 µm Ibidi layer remaining on top of the resin block was removed with a 20° diamond trimming knife (Diatome, Switzerland). Serial sections (300 nm) were cut on an ultramicrotome (Leica EM UC7) using a 35° diamond knife (Diatome, Switzerland) and collected onto 50 mesh copper grids (Agar Scientific) coated with 1% formvar in chloroform. Sections (300 nm) were post-stained with 2% uranyl acetate in 70% methanol for 5 min at room temperature, followed by Reynolds’ lead citrate for 2 min at room temperature. Stained sections were screened for regions of interest using a JEM-2100+ transmission electron microscope (JEOL) operated at 200 kV, controlled with SerialEM. Tomographic tilt series of selected regions were subsequently acquired on a Tecnai F30 (Thermo Fisher) microscope at 300 kV. Three-dimensional reconstructions were generated using eTOMO.

### Fate-resolved sister-cell single cell genomic sequencing and analysis

#### I. Micronucleus-fate-matched sister-cell isolation

BrdU (40µM final concentration) was added to hTERT RPE-1 cells expressing H2B-mKeima. 16 hours later cells were arrested in mitosis with 100ng/ml nocodazole for 6 hours, followed by mitotic shake off (as described above) into an Ibidi chamber slide (Ibidi, 80806) in complete medium with BrdU. Within 1 hour of shake-off slides were transferred to a pre-heated LSM 900 confocal microscope (Zeiss) (Plan-Apochromat ×20/0.8 M27 air objective, 37 degrees with CO_2_) and large tiled images taken at 20 minute intervals for 3 hours. Cells were then briefly washed with pre-heated medium with BrdU to remove unattached cells and imaging of the same region resumed for a further 12-15 hours. The photoactivatable dye DACT-1^6^ (10µM final concentration) and, optionally, lysosome inhibitor Bafilomycin A* (200nM final concentration, to prevent additional chromophagy events) were then added to the slide without disrupting the imaging, and imaging continued until regions of interest had been defined and updated to live positions using the experimental regions tab in ZEN. While imaging was ongoing the fate of micronucleated cells and identity of corresponding sister cells was determined by manual assessment of the timecourse imaging files up to the point of DACT-1 and Baf A addition. Once all ROI were defined (corresponding to sister-cell pairs with the desired micronucleus-fate) and updated to live positions, photoactivation of DACT-1 was carried out using a 405nm laser. Hoechst 33342 (5 µg/ml final concentration) was then added and cells incubated for 30 minutes, trypsinized, and resuspended in 8% FBS in 1× PBS, supplemented with 200nM Bafilomycin A. Single photoactivated cells were then sorted on a BD FACSAria or BDFACS Discover S8 using a 100 µm nozzle into lysis buffer (recipe: 2.4 ml PBS (no Ca & Mg), 2.125ml Profreeze, 375 ul DMSO 100 ul 10% Np40 substitute) in a flat-bottomed 96-well plate, spun down, and then transferred to a −70°C freezer. *\* added to all experiments except 4 pairs of each acidified / non-acidified with no discernable impact*.

#### II. Strand-seq

Single cell template strand genomic sequencing (Strand-seq) libraries were prepared using a liquid handling robotic platform as described previously^50^. Libraries were sequenced on an AVITI Medium 75bp PE (2×75), followed by demultiplexing. Reads were aligned to GRCh38 reference assembly with BWA-MEM v.0.7.17 and MosaiCatcher v2^71^, and further processed as described below.

#### III. Sister-cell identification

Sister cell pairs were identified computationally from the sequenced Strand-seq libraries, using the strandtools computational framework^6^. The categorical strand-state annotation assigned to each genomic bin was first transformed into a numerical representation, with WW states encoded as 1, WC states as 0, and CC states as −1. Based on these encoded strand-state profiles, pairwise Pearson correlation coefficients were calculated across all single cells within each experiment. Cell pairs exhibiting a correlation coefficient < −0.5 were visually inspected and classified as sister cells.

#### IV. Identification of de novo chromosomal loss events from Strand-seq from sister cells

Strand-seq uses BrdU-mediated marking of newly-synthesized strands during DNA replication, to distinguish them from template strands. Each replicated chromosome therefore produces two sister chromatids, one in W orientation and one in C orientation, each with one template originating from unwinding of the original double-helix. This semi-conservative property of DNA replication essentially grants that W and C template strands among sisters are always present in equal numbers, regardless of copy number changes and in absence of chromosome degradation. Building on this model, we pooled and summed W and C reads across sister cell pairs, for each chromosome arm and generated a scatterplot. If chromosome integrity is maintained, W and C template strands display a balanced distribution with an approximate 1:1 ratio (scenario 1) across each sister chromosome pair. Minor deviations from this balance may occur following chromosome missegregation events, due to differences in library composition and normalization. In contrast, *de novo* chromosome loss, for example when DNA becomes sequestered in an acidified micronucleus and subsequently degraded, would result in the loss of one template strand. This would skew the W ratio toward the 1:2 or 2:1 (scenario 2) in disomic regions. Importantly, such skewing would affect only the chromosomes experiencing chromosomal loss, while other chromosomes within the same cell would remain balanced and represent an internal control. *De novo* CAs were identified by chromosome arm W ratio changes, and by automated copy number calling using strandtools, as described previously^6^.

### Impact of micronucleus fates on interphase chromosome fragmentation

hTERT RPE-1 cells expressing H2B-mKeima and HaloTag-NLS were imaged for 18 hours following release from a 6 hour nocodazole arrest. Cells were then stained for yH2AX and live-imaging fates and yH2AX intensities (mean relative to primary nuclei) determined for each micronucleus.

### Impact of chromophagy on the intergenerational transmission of micronuclei derived chromosomal material

hTERT RPE-1 cells expressing H2B-mKeima were transfected with siRNA against p53 18 hours before a 6 hour nocodazole arrest to facilitate cell cycle progression after sustained mitotic arrest and aneuploidy induction. After mitotic shake-off, cells were imaged at 20 minute intervals for 42h on a Zeiss Cell discoverer 7 so that most of the cells were in their second interphase. Cells were then fixed, stained for yH2AX, and live-imaging recordings were used to determine the presence and fate of micronuclei in generation 1, and to determine the location of generation 2 daughter cells. Daughter cells were scored for the presence of Micronuclei (MN), yH2AX+ve MN, and MN bodies (identified as large yH2AX territories within the main nucleus).

### TUNEL, EdU and Hoechst staining for analysis of DNA degradation

For the analysis of DNA degradation following MN degradation, hTERT RPE-1 H2B-mKeima were live imaged for 4 hours following nocodazole shake-off and then fixed; the live-imaging was used to determine MN degradation and its timing before fixation for subsequent analysis.

For TUNEL staining, the CF640R TUNEL Assay Kit (Biotium, 30074) was used following the manufacturer’s protocol. Briefly, cells were permeabilized with 0,2% Triton X-100 for 20 minutes, washed with equilibration buffer for 5 minutes and then incubated for 60 min at 37C with TdT enzyme to allow staining.

For EdU staining the Click-iT™EdU Imaging kit was used (Thermofisher, C10338). Cells were incubated for 24 hours before nocodazole shake-off with 2mM EdU to ensure complete DNA labelling and after fixation the manufacturer’s protocol was followed for EdU staining.

After staining with either the TUNEL or EdU kit, cells were incubated with Hoechst for 10 minutes to stain DNA in MN. TUNEL, EdU and Hoechst intensities in each MN were normalized to the intensities of the same marker in the respective primary nucleus (after background subtraction).

## References

1. Lengauer, C., Kinzler, K. W. & Vogelstein, B. Genetic instabilities in human cancers. Nature 396, 643–649 (1998).

2. Cosenza, M. R., Rodriguez-Martin, B. & Korbel, J. O. Structural Variation in Cancer: Role, Prevalence, and Mechanisms. Annu Rev Genomics Hum Genet 23, 123–152 (2022).

3. Sansregret, L., Vanhaesebroeck, B. & Swanton, C. Determinants and clinical implications of chromosomal instability in cancer. Nature Reviews Clinical Oncology 15, 139–150 (2018).

4. He, B. et al. Chromosomes missegregated into micronuclei contribute to chromosomal instability by missegregating at the next division. Oncotarget 10, 2660–2674 (2019).

5. Umbreit, N. T. et al. Mechanisms generating cancer genome complexity from a single cell division error. Science (2020) doi:10.1126/science.aba0712.

6. Cosenza, M. R. et al. Origins of chromosome instability unveiled by coupled imaging and genomics. Nature (2025) doi:10.1038/s41586-025-09632-5.

7. Zhang, C.-Z. et al. Chromothripsis from DNA damage in micronuclei. Nature 522, 179–184 (2015).

8. Stephens, P. J. et al. Massive genomic rearrangement acquired in a single catastrophic event during cancer development. Cell 144, 27–40 (2011).

9. Rausch, T. et al. Genome sequencing of pediatric medulloblastoma links catastrophic DNA rearrangements with TP53 mutations. Cell 148, 59–71 (2012).

10. Hanahan, D. & Weinberg, R. A. Hallmarks of cancer: the next generation. Cell 144, 646–674 (2011).

11. A Mazzagatti, J. L. Engel, P. Ly. Boveri and beyond: Chromothripsis and genomic instability from mitotic errors. Molecular Cell 84, 55–69 (2024).

12. Hatch, E. M., Fischer, A. H., Deerinck, T. J. & Hetzer, M. W. Catastrophic nuclear envelope collapse in cancer cell micronuclei. Cell 154, 47–60 (2013).

13. Liu, S. et al. Nuclear envelope assembly defects link mitotic errors to chromothripsis. Nature 561, 551–555 (2018).

14. Lin, Y.-F. et al. Mitotic clustering of pulverized chromosomes from micronuclei. Nature 618, 1041–1048 (2023).

15. Shoshani, O. et al. Chromothripsis drives the evolution of gene amplification in cancer. Nature 591, (2021).

16. Zhang, C.-Z., Leibowitz, M. L. & Pellman, D. Chromothripsis and beyond: rapid genome evolution from complex chromosomal rearrangements. Genes Dev. 27, 2513–2530 (2013).

17. Krupina, K., Goginashvili, A. & Cleveland, D. W. Scrambling the genome in cancer: causes and consequences of complex chromosome rearrangements. Nature Reviews Genetics 25, 196–210 (2023).

18. Dikic, I. & Elazar, Z. Mechanism and medical implications of mammalian autophagy. Nature Reviews Molecular Cell Biology 19, 349–364 (2018).

19. Rello-Varona, S. et al. Autophagic removal of micronuclei. Cell Cycle 11, 170–176 (2012).

20. Martin, S. et al. A p62-dependent rheostat dictates micronuclei catastrophe and chromosome rearrangements. Science 385, eadj7446 (2024).

21. Holdgaard, S. G. et al. Selective autophagy maintains centrosome integrity and accurate mitosis by turnover of centriolar satellites. Nat Commun 10, 4176 (2019).

22. Liu, E. Y. et al. Loss of autophagy causes a synthetic lethal deficiency in DNA repair. Proceedings of the National Academy of Sciences 112, 773–778 (2015).

23. Marom, S. et al. Aberrant inheritance of extrachromosomal DNA amplifications promotes cancer evolution. bioRxiv 2025.09.19.677276 (2025) doi:10.1101/2025.09.19.677276.

24. The ICGC/TCGA Pan-Cancer Analysis of Whole Genomes Consortium. Pan-cancer analysis of whole genomes. Nature 578, 82–93 (2020).

25. Duijf, P. H. G., Schultz, N. & Benezra, R. Cancer cells preferentially lose small chromosomes. Int J Cancer 132, 2316–2326 (2013).

26. H. Katayama, T. Kogure, N. Mizushima, T. Yoshimori, A. Miyawaki. A Sensitive and Quantitative Technique for Detecting Autophagic Events Based on Lysosomal Delivery. Chemistry & Biology 18, 1042–1052 (2011).

27. Zasadil, L. M. et al. Cytotoxicity of paclitaxel in breast cancer is due to chromosome missegregation on multipolar spindles. Sci Transl Med 6, 229ra43 (2014).

28. Khaminets, A. et al. Regulation of endoplasmic reticulum turnover by selective autophagy. Nature 522, 354–358 (2015).

29. Jaber, N. et al. Class III PI3K Vps34 plays an essential role in autophagy and in heart and liver function. Proceedings of the National Academy of Sciences 109, 2003–2008 (2012).

30. Dowdle, W. E. et al. Selective VPS34 inhibitor blocks autophagy and uncovers a role for NCOA4 in ferritin degradation and iron homeostasis in vivo. Nat Cell Biol 16, 1069–1079 (2014).

31. Kwon, M., Leibowitz, M. L. & Lee, J.-H. Small but mighty: the causes and consequences of micronucleus rupture. Experimental & Molecular Medicine 52, 1777–1786 (2020).

32. Maciejowski, J., Li, Y., Bosco, N., Campbell, P. J. & de Lange, T. Chromothripsis and Kataegis Induced by Telomere Crisis. Cell 163, 1641–1654 (2015).

33. Ly, P. et al. Chromosome Segregation Errors Generate a Diverse Spectrum of Simple and Complex Genomic Rearrangements. Nature genetics 51, 705 (2019).

34. Trivedi, P., Steele, C. D., Au, F. K. C., Alexandrov, L. B. & Cleveland, D. W. Mitotic tethering enables inheritance of shattered micronuclear chromosomes. Nature 618, 1049–1056 (2023).

35. Young, A. M., Gunn, A. L. & Hatch, E. M. BAF facilitates interphase nuclear membrane repair through recruitment of nuclear transmembrane proteins. Mol. Biol. Cell 31, 1551–1560 (2020).

36. Gatica, D., Lahiri, V. & Klionsky, D. J. Cargo recognition and degradation by selective autophagy. Nat Cell Biol 20, 233–242 (2018).

37. de Castro, I. J., Gil, R. S., Ligammari, L., Di Giacinto, M. L. & Vagnarelli, P. CDK1 and PLK1 coordinate the disassembly and reassembly of the nuclear envelope in vertebrate mitosis. Oncotarget 9, 7763–7773 (2018).

38. Orr, B. et al. An anaphase surveillance mechanism prevents micronuclei formation from frequent chromosome segregation errors. Cell Rep 37, 109783 (2021).

39. Samwer, M. et al. DNA Cross-Bridging Shapes a Single Nucleus from a Set of Mitotic Chromosomes. Cell 170, 956–972.e23 (2017).

40. Lee, K. K. et al. Distinct functional domains in emerin bind lamin A and DNA-bridging protein BAF. J Cell Sci 114, 4567–4573 (2001).

41. Jamin, A. & Wiebe, M. S. Barrier to Autointegration Factor (BANF1): interwoven roles in nuclear structure, genome integrity, innate immunity, stress responses and progeria. Curr Opin Cell Biol 34, 61–68 (2015).

42. Shumaker, D. K., Lee, K. K., Tanhehco, Y. C., Craigie, R. & Wilson, K. L. LAP2 binds to BAF.DNA complexes: requirement for the LEM domain and modulation by variable regions. EMBO J 20, 1754–1764 (2001).

43. Lewis, R. et al. LBR and LAP2 mediate heterochromatin tethering to the nuclear periphery to preserve genome homeostasis. Nat Cell Biol (2026) doi:10.1038/s41556-025-01822-7.

44. Chang, L. et al. Nuclear peripheral chromatin-lamin B1 interaction is required for global integrity of chromatin architecture and dynamics in human cells. Protein Cell 13, 258–280 (2022).

45. Molitor, T. P. & Traktman, P. Depletion of the protein kinase VRK1 disrupts nuclear envelope morphology and leads to BAF retention on mitotic chromosomes. Mol Biol Cell 25, 891–903 (2014).

46. Paouneskou, D. et al. BAF-1–VRK-1 mediated release of meiotic chromosomes from the nuclear periphery is important for genome integrity. Nature Communications 16, 10446 (2025).

47. Serafim, R. A. M. et al. Development of Pyridine-based Inhibitors for the Human Vaccinia-related Kinases 1 and 2. ACS Med Chem Lett 10, 1266–1271 (2019).

48. Schreiner, S. M., Koo, P. K., Zhao, Y., Mochrie, S. G. J. & King, M. C. The tethering of chromatin to the nuclear envelope supports nuclear mechanics. Nat Commun 6, 7159 (2015).

49. Falconer, E. et al. DNA template strand sequencing of single-cells maps genomic rearrangements at high resolution. Nature methods 9, (2012).

50. Sanders, A. D. et al. Single-cell analysis of structural variations and complex rearrangements with tri-channel processing. Nature biotechnology 38, (2020).

51. Worrall, J. T. et al. Non-random Mis-segregation of Human Chromosomes. Cell Rep 23, 3366–3380 (2018).

52. Watson, J. D. & Crick, F. H. C. Molecular Structure of Nucleic Acids: A Structure for Deoxyribose Nucleic Acid. Nature 171, 737–738 (1953).

53. Papathanasiou, S. et al. Heritable transcriptional defects from aberrations of nuclear architecture. Nature 619, 184–192 (2023).

54. Schäfer, P., Cymerman, I. A., Bujnicki, J. M. & Meiss, G. Human lysosomal DNase IIα contains two requisite PLD-signature (HxK) motifs: Evidence for a pseudodimeric structure of the active enzyme species. Protein Science : A Publication of the Protein Society 16, 82 (2007).

55. Lan, Y. Y., Londoño, D., Bouley, R., Rooney, M. S. & Hacohen, N. Dnase2a deficiency uncovers lysosomal clearance of damaged nuclear DNA via autophagy. Cell Rep 9, 180–192 (2014).

56. Papini, D., Levasseur, M. D. & Higgins, J. M. G. The Aurora B gradient sustains kinetochore stability in anaphase. Cell Rep 37, 109818 (2021).

57. Sen, O., Harrison, J. U., Burroughs, N. J. & McAinsh, A. D. Kinetochore life histories reveal an Aurora-B-dependent error correction mechanism in anaphase. Dev Cell 56, 3082–3099.e5 (2021).

58. Galluzzi, L. et al. Autophagy in malignant transformation and cancer progression. The EMBO Journal (2015) doi:10.15252/embj.201490784.

59. Hewitt, G. & Korolchuk, V. I. Repair, Reuse, Recycle: The Expanding Role of Autophagy in Genome Maintenance. Trends Cell Biol 27, 340–351 (2017).

60. Lascaux, P. et al. TEX264 drives selective autophagy of DNA lesions to promote DNA repair and cell survival. Cell 187, 5698–5718.e26 (2024).

61. Dou, Z. et al. Autophagy mediates degradation of nuclear lamina. Nature 527, 105–109 (2015).

62. Nassour, J. et al. Autophagic cell death restricts chromosomal instability during replicative crisis. Nature 565, 659–663 (2019).

63. Baylin, S. B. & Jones, P. A. Epigenetic Determinants of Cancer. Cold Spring Harb Perspect Biol 8, a019505 (2016).

64. Flavahan, W. A., Gaskell, E. & Bernstein, B. E. Epigenetic plasticity and the hallmarks of cancer. Science (2017) doi:10.1126/science.aal2380.

65. Feinberg, A. P., Koldobskiy, M. A. & Göndör, A. Epigenetic modulators, modifiers and mediators in cancer aetiology and progression. Nature Reviews Genetics 17, 284–299 (2016).

66. Santana-Codina, N., Mancias, J. D. & Kimmelman, A. C. The Role of Autophagy in Cancer. Annu Rev Cancer Biol 1, 19–39 (2017).

67. Cahill, D. P., Kinzler, K. W., Vogelstein, B. & Lengauer, C. Genetic instability and darwinian selection in tumours. Trends Cell Biol 9, M57–60 (1999).

68. Weaver, B. A. A., Silk, A. D., Montagna, C., Verdier-Pinard, P. & Cleveland, D. W. Aneuploidy acts both oncogenically and as a tumor suppressor. Cancer Cell 11, 25–36 (2007).

69. Silk, A. D. et al. Chromosome missegregation rate predicts whether aneuploidy will promote or suppress tumors. Proc Natl Acad Sci U S A 110, E4134–41 (2013).

70. Ershov, D. et al. TrackMate 7: integrating state-of-the-art segmentation algorithms into tracking pipelines. Nat Methods 19, 829–832 (2022).

71. Weber, T., Cosenza, M. R. & Korbel, J. MosaiCatcher v2: a single-cell structural variations detection and analysis reference framework based on Strand-seq. Bioinformatics 39, btad633 (2023).

