## Supplemental figures for "Selective autophagy of whole micronuclei suppresses chromosomal instability"

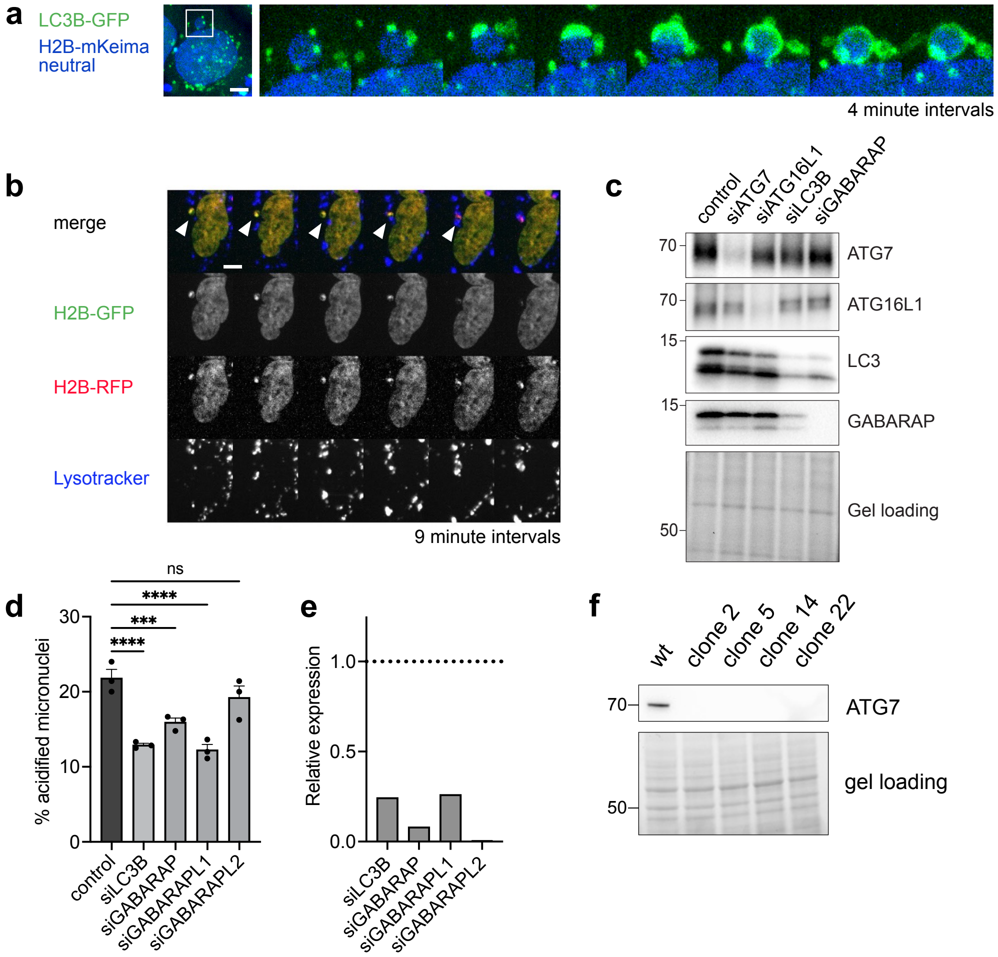


**Figure S1**

**a,** Step-wise capture of a large micronucleus within an LC3B positive autophagosome (scale bar = 5µm). **b,** MN acidification upon lysosome fusion using H2B-GFP-RFP reporter (scale bar = 5µm). **c,** siRNA knockdown of autophagy pathway proteins by western blot (for Fig 1i). **d,** MN acidification rate upon knockdown of the different ATG8s expressed in hTERT RPE-1 (n≥100MN per experiment, 3 separate experiments). **e,** qPCR result showing knockdown of the target ATG8s. **f,** Western blot showing knock-out of ATG7 in the selected clones.

Data are shown as mean ± sem, and were analysed using a one-way ANOVA (d) (ns = non-significant, *** = p ≤ 0.001, **** = p ≤ 0.0001)

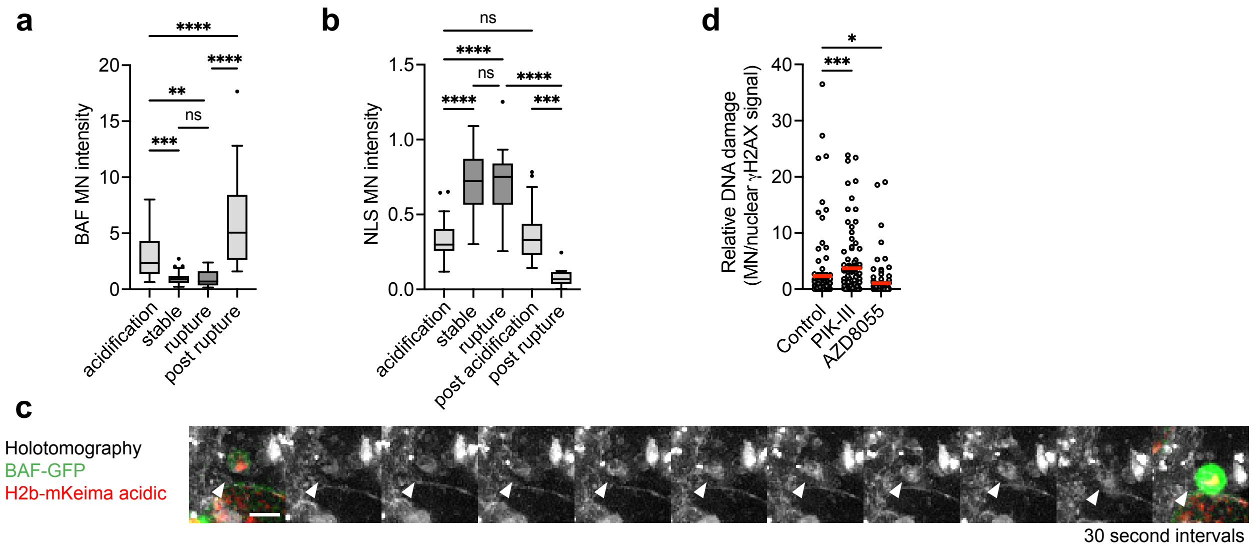


**Figure S2**

**a,** BAF levels on MN, normalized relatively to the main nucleus, based on the different MN fates. (n between 18 and 34 individual MN, 3 separate experiments). **b,** NLS levels on MN, normalized relatively to the main nucleus, based on the different MN fates (n between 13 and 26 individual MN, 2 separate experiments). **c,** Holotomography montage showing absence of membrane recruitment around MN before rupture of the membrane, shown by sudden BAF over recruitment (scale bar = 2µm). **d,** Quantification of MN DNA damage (relative to the primary nucleus) after autophagy inhibition (PIK-III) or induction (AZD8055) (n between 93 and 106 individual MN, 3 separate experiments).

Data are shown as mean ± sem, and were analysed using a one-way ANOVA (a, b, d) (ns = non-significant, * = p ≤ 0.05, ** = p ≤ 0.01, *** = p ≤ 0.001, **** = p ≤ 0.0001)

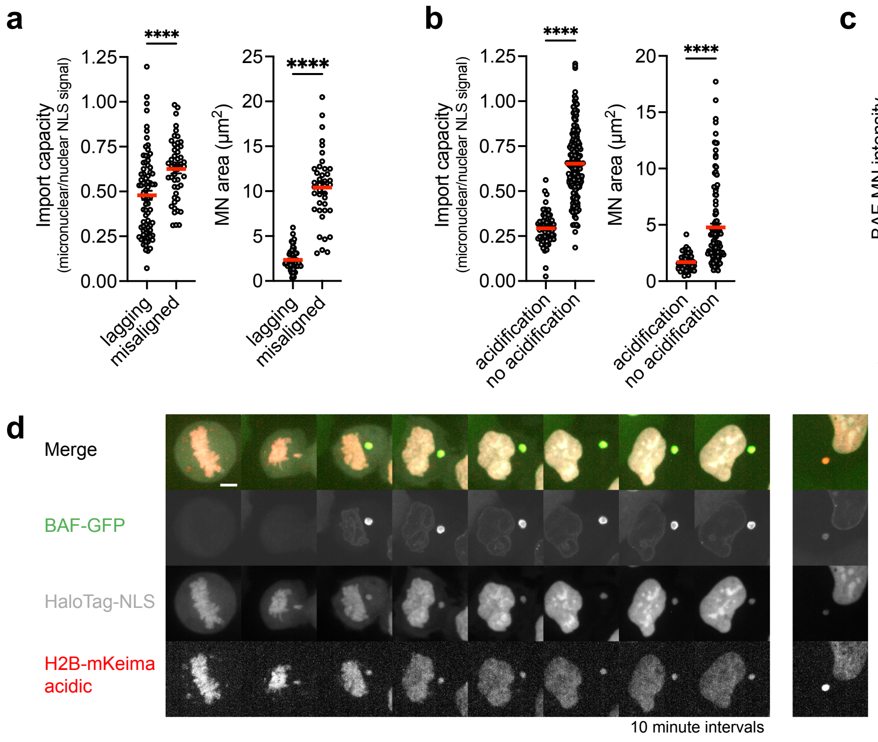


**Figure S3**

**a,** Import capacity (left) and MN area (right) of lagging and misaligned MN following GSK+NMP induction (n=80 and 49 MN from n=3 experiments). **b,** Import capacity (left) and MN area (right) of MN that undergo acidification or not (n=46 and 116 MN from n=3 experiments). **c,** MN levels of BAF (left) (n=12 and 29 MN from n=3 experiments), Lamin B1 (centre) (n=32 and 42 MN from n=3 experiments), and Emerin (right) (n=15 and 25 MN from n=2 experiments) of MN that undergo acidification or not. **d,** BAF over recruitment at mitotic exit on lagging MN that later undergoes acidification (scale bar = 5μm).
Data are shown as mean ± sem, and were analysed using an unpaired t-test (a, b, c) (* = p ≤ 0.05,**** = p ≤ 0.0001)

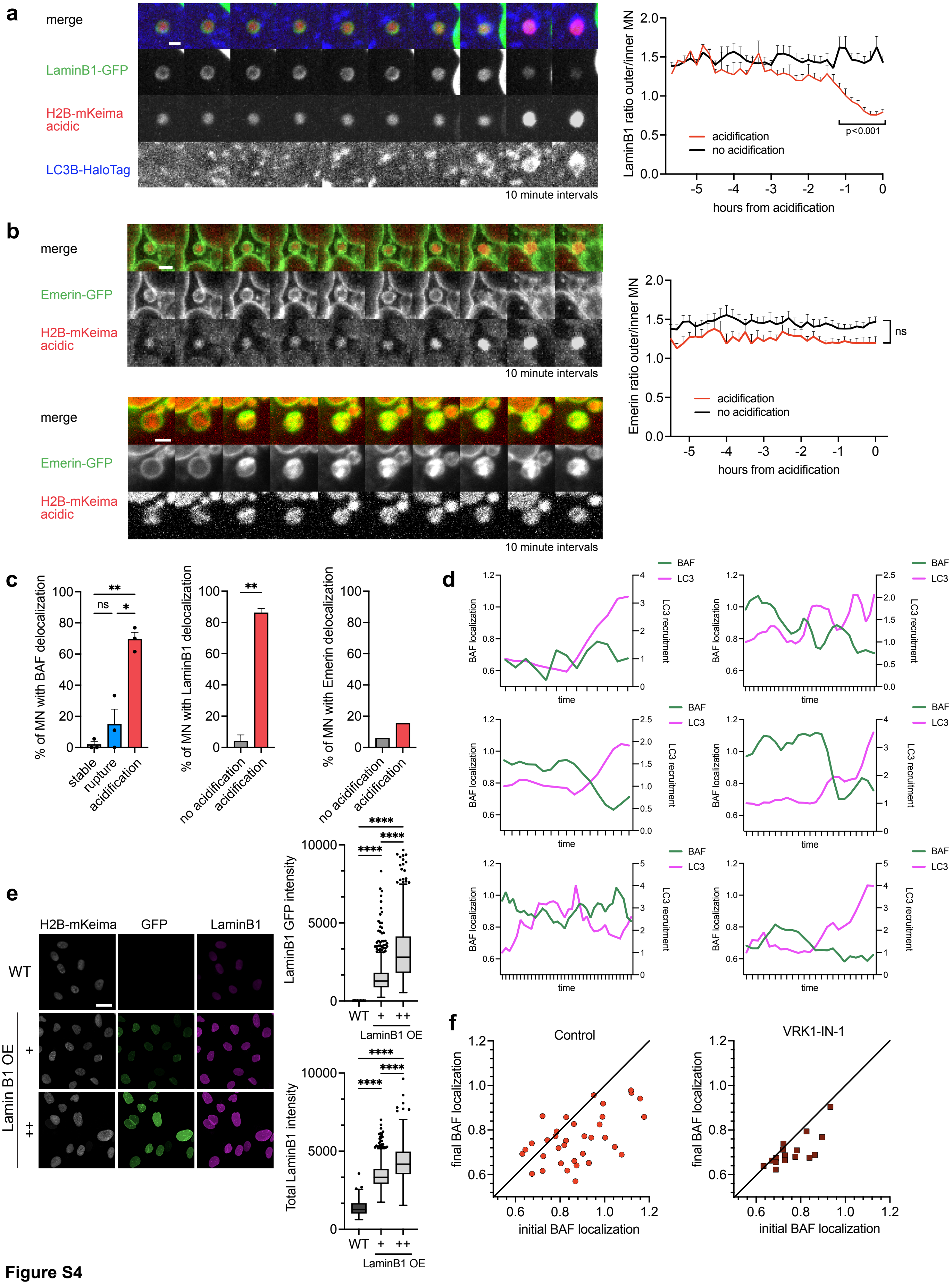


**Figure S4**

**a,** Montage showing Lamin B1 dissociation from the nuclear membrane before MN acidification (left) and timecourse of localisation by fate (right) (scale bar = 2μm). **b,** Montage showing Emerin localisation before MN acidification (top), Emerin infiltration upon MN rupture (bottom) and localisation timecourse by fate (right) (scale bar = 2μm). **c,** Quantification of MN showing sustained dissociation of BAF (left), Lamin B1 (centre) or Emerin (right) based on the different MN fates. **d,** Representative plots of single MN showing the dynamics of LC3 and BAF before acidification (from 10 minute interval imaging). **e,** Representative images (top) and quantification (bottom) of GFP-Lamin B1 and total Lamin B1 in the analysed cell lines (scale bar = 20μm). **f,** Plots showing initial and final BAF localization of individual MN undergoing acidification in basal condition (left) or VRK1-IN-1 treated cells (right).

Data are shown as mean ± sem, and were analysed using a two-way ANOVA (a, b), one-way ANOVA (c-left, e), unpaired t-test (c-centre) (* = p ≤ 0.05,**** = p ≤ 0.0001)

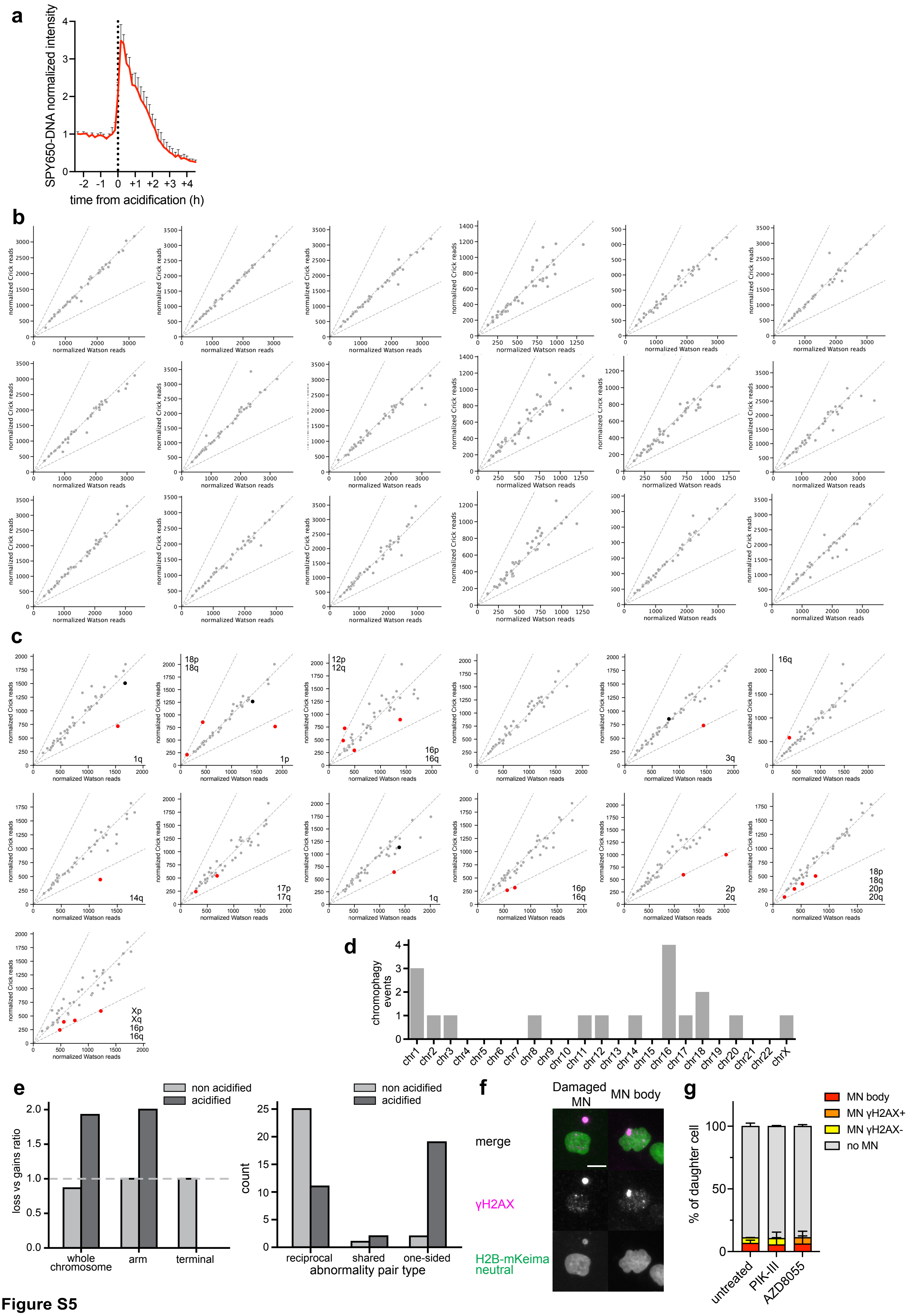


**Figure S5**

**a,** Timecourse of SPY650 DNA dye signal intensity in live-cells undergoing acidification. **b,** W:C ratio plots of 18 additional sister cell pairs with live-imaging verified non-acidification fates. **c,** W:C ratio plots of 13 additional sister cell pairs with live-imaging verified acidification fates (chromophagy affected chromosome arms are highlighted in red and indicated in the graph while corresponding unaffected arms are in black). **d,** Chromosomal distribution of verified chromophagy events. **e,** Loss vs gain ratio by copy number variation type (left) and count of chromosomal abnormalities by sister cell reciprocity status (right) for cells with acidified and not acidified MN. **f,** Representative images of daughters of micronucleated parental cells with γH2AX+ micronuclei (left), or MN body (right), characteristic of intergenerational transmission of fragmented (micronuclei derived) chromosomal material. **g,** MN and MN bodies in the daughters of non-micronucleated parental cells following pharmacological autophagy inhibition (PIK-III) or stimulation (AZD8055) (scale bar = 10μm).
